# Differential induction of *Medicago truncatula* defence metabolites in response to rhizobial symbiosis and pea aphid infestation

**DOI:** 10.1101/2024.08.14.607928

**Authors:** Goodluck Benjamin, Marie Pacoud, Stéphanie Boutet, Gilles Clement, Renaud Brouquisse, Jean-Luc Gatti, Marylène Poirié, Pierre Frendo

**Author notes:** Both authors supervised this work (Marylène Poirié, Pierre Frendo).

## Abstract

- Legumes symbiosis with rhizobial nitrogen-fixing bacteria enable them to grow in nitrate-depleted soils. Rhizobial symbioses also induces systemic plant defence against bioagressors.
- We investigate how nitrogen-fixing symbiosis (NFS) in the legume *Medicago truncatula* can prime plant defence against the pea aphid *Acyrthosiphon pisum*. We analysed metabolite modification both by LC-MS and GC-MS and defence pathway gene expression by qPCR in leaves of both NFS and nitrate-fed (non-inoculated; NI) plants after aphid infestation (Amp).
- The accumulation of primary and secondary metabolites was modulated by both NFS and aphid infestation. 62 defense-related metabolites such as salicylate, pipecolate, gentisic acid and several soluble sugars were differentially regulated by aphid infestation in both NFS and NI conditions. 19 metabolites, including triterpenoid saponins, accumulated specifically in NFS_Amp conditions. Gene expression analysis showed that aphid-infested plants exhibited significantly higher expression of *Chalcone isomerase*, *flavonol synthase*, *hydroxyisoflavone-O-methyl transferase* and *Pterocarpan synthase*, while *D-pinitol dehydrogenase* was only significantly induced in NI infested leaves.
- Our data suggest that NFS, in addition to being a plant nitrogen provider, stimulates specific legume defenses upon pest attack and should also be considered as a potential tool in Integrated Pest Management strategy.

## Introduction

Plants are under constant threat from pathogens and insect pests such as sap-feeding aphids (National Research Council *et al*., 1985). Aphids are deleterious agronomic pests, not only because they feed on phloem sap and therefore weaken the plant, but also because they are vectors for various plant viruses (Ng & Perry, 2004). More than 5000 species of aphids are known today, a diversity that is partly due to sympatric speciation initiated by individuals adapting to new host plants (Diehl & Bush, 1984; Drès & Mallet, 2002). During feeding, the aphid stylet injures plant cells, inject saliva, and sucks up tiny amounts of the cell content to decide on plant acceptance (Martin *et al*., 1997; Lu *et al*., 2016). Previous studies have revealed that the secreted proteins present in saliva trigger plant responses (Pitino & Hogenhout, 2013; Rodriguez & Bos, 2013). These saliva proteins act as herbivore-associated molecular patterns (HAMPs) (Kaloshian & Walling, 2016) that bind to host plant pattern recognition receptors (PRRs) and trigger an immune response associated with the production of reactive oxygen species (ROS) and defence hormones Salicylic Acid (SA) and Jasmonic Acid (JA) pathways (Wu & Baldwin, 2010; Herrera-Vásquez *et al*., 2015). The increase of SA and JA regulate the accumulation of various primary and secondary metabolites that play a role in plant defence (Isah, 2019) as feeding deterrents or toxins that decrease food intake or food use efficiency, decrease survival and reproduction of the pest, or indirectly act as attractants for its natural enemies (War *et al*., 2012).The secondary metabolites involved in this process include terpenoids, phenolics, cyanogenic glycosides, glucosinolates, and alkaloids (Maag *et al*., 2015; Züst & Agrawal, 2016). For example, high levels of saponins and phenolic compounds increase aphid mortality and result in reduced pest population (Züst & Agrawal, 2016), flavonoid glycosides also reduce aphid fecundity, while nitrogen-containing compounds cause host plant rejection (Kordan *et al*., 2012).

One property of legume plants is their ability to associate with rhizobia to perform nitrogen fixation symbiosis (NFS), i.e., the conversion of atmospheric N_2_ into plant-usable ammonium. The microbial partner supplies assimilable nitrogen to the plant in exchange for carbon resources and a protective environment in the root nodules (Lee & Hirsch, 2006). NFS has also been reported to potentially provide defence priming to plants against bio-aggressors (Benjamin *et al*., 2022). For example, in pea (*Pisum sativum*), rhizobium inoculation decreased *Didymella pinodes* disease severity and significantly reduced the seed infection level (Desalegn *et al*., 2016; Ranjbar Sistani *et al*., 2017). NFS has also been reported to modulate resistance to biotrophic pathogens in both *Medicago truncatula* and pea by reducing penetration and sporulation of the powdery mildew fungus *Erysiphe pisi* (Smigielski *et al*., 2019).

We have previously demonstrated that NFS also influences the *Medicago truncatula*–pea aphid interaction and general plant defence response (Pandharikar *et al*., 2020). A detrimental effect of rhizobia-inoculated plants on aphid development was observed, with lower adult weights compared to aphids from nitrate-fed plants. *Pathogenesis Related 1* (*PR1*) gene expression was upregulated in aphid-infested shoots, indicating the activation of SA-dependent defence. Moreover, a significantly higher expression of *Proteinase Inhibitor (PI)* gene, a marker for the JA transduction pathway, was observed in NFS plants compared to nitrate-fed plants (Pandharikar *et al*., 2020).

Due to the observations from our preliminary studies, we have analysed by metabolomics, the impact of the nitrogen source (KNO_3_ *vs* NFS) on *M. truncatula* infestation by pea aphid. To this end we analysed leaf metabolite profiles of NFS plants with and without aphid infestation and compared them to KNO_3_-fed plants (Non Inoculated; NI) and to obtain a maximum coverage of the leaf metabolites, untargeted metabolomic was done both by gas chromatography/mass spectrometry (GC-MS) and liquid chromatography/mass spectrometry (LC-MS). Our results showed that both primary and secondary metabolisms are significantly modulated by nitrogen source and aphid infestation. Gene expression analysis of enzymes involved in the secondary metabolite synthesis pathways showed that regulation of secondary metabolism is partially mediated by the modulation of gene expression. We identified specific metabolites which are differentially regulated in NFS plants compared to NI plants under aphid infestation.

## Materials and Methods

### Biological materials and experimental design

The pea aphid *Acyrtosiphon pisum* clone, YR2-amp, further named YR2, is derived from a clover biotype line collected in England that was freed from the secondary symbiont *Regiella insecticola* by ampicillin treatment (Simon *et al*., 2011). This aphid line is stable (more than 15 years old) and was maintained on fava bean, 20°C, 16:8h light/dark cycle. To synchronize aphid, 20-40 apterous female adults were placed in a petri dish containing a fava bean leaf and let to reproduce for 24hrs. Then, 10 nymphs (L1) were collected and used for infestation.

For one experiment, four batches of 5 pots containing each six *Medicago truncatula* A17 plants were grown as previously described (Pandharikar *et al*., 2020) (Fig. S1). After 12 days, two groups were inoculated with the nitrogen-fixing bacteria *Sinorhizobium meliloti* 2011 (nitrogen-fixing symbiosis, NFS condition), and two were supplemented once with 10 ml of 5 mM KNO_3_ solution (non-inoculated, NI condition). Seven days after (the time to NFS plants to develop nodules), for both NFS and NI conditions, one batch of NI plants and one batch of NFS plants were infested with aphids (10 L1/pots). The aphid nymphs were then let to develop into adult during 12 days. All pots were individually isolated in a ventilated plastic box and maintained at 20℃ under 16:8h light/dark photoperiod. For each condition, control pots were treated and maintained under the same conditions except that no aphids were introduced in the plastic box. At the end of the 12 days, harvested leaves were used immediately or frozen in liquid nitrogen and stored at -80°C.

Rhizobium inoculation was done with a streptomycin-resistant strain of *Sinorhizobium meliloti* 2011. It was cultured on Luria-Bertani medium supplemented with 2.5mM CaCl_2_ and MgSO_4_ (LBMC) and streptomycin at 200μg ml^-1^ for 3 days at 30°C, then transferred and grown in LBMC liquid medium for 24h, pelleted at 5,000g, washed twice with sterile distilled water, and resuspended in sterile distilled water to a final optical density of 0.05 (OD_600_). Each NFS plant was inoculated with 10 mL of this *S. meliloti* suspension.

A total of 4 biological replicates (i.e. 4 times 4 pots of six plants for each condition), produced as described above were used for analysis carried out in this study.

Analysis of metabolites using gas chromatography/mass spectrometry (GC-MS) Leaves were ground in liquid nitrogen to obtain a fine powder and 50 mg was resuspended in 1ml of cold (-20°C) Water:Acetonitrile:Isopropanol (2:3:3 in volume) containing Ribitol 4μg/ml as internal standard. After extraction under shaking (10 min, 4°C), insoluble material was removed by centrifugation at 20,000g for 5min and 50μl were collected and dried overnight (SpeedVac™) and used immediately or stored at -80°C. Three blank tubes underwent the same treatments to estimate possible contamination. A quality control was made by pooling an equal volume of each condition.

Samples were warmed 15 min before opening and dried again for 1.5h at 35°C before addition of 10μl of 20 mg ml^-1^ methoxyamine in pyridine and the reaction was performed for 90 min at 28°C under continuous shaking. 90μl of N-methyl-N-trimethylsilyltrifluoroacetamide (MSTFA) were then added and the reaction continued for 30 min at 37°C. After cooling, 45μl were taken for injection. 1μl of derivatized sample was injected in splitless and split (1:30) modes on a gas chromatograph (Agilent 7890A; Santa Clara, USA) coupled to a mass spectrometer (Agilent 5977B) with a heated separation column (Rxi-5SilMS; Restek, Lisses, France) (temperature ramp: 70°C for 7 min then 10°C/min to 330°C for 5 min; run length 38 min). Helium flow was constant at 0.7 mL/min. 5 scans per second were acquired spanning a 50 to 600Da range. Instrument was tuned with PFTBA with the *m/z* 69 and *m/z* 219 of equal intensities. Samples were randomized. Three independent quality controls were injected at the beginning, middle and end of the analysis for monitoring the derivatization stability. An alkane mix (C10, C12, C15, C19, C22, C28, C32, C36) was injected during the run for external calibration. Three independent derivatizations of the quality control were injected at the beginning, in the middle and at the end of the series. A response coefficient was determined for 4ng each of a set of 103 metabolites, respectively to the same amount of ribitol. This compound was used to give an estimation of the absolute concentration of the metabolite in what we may call a “one point calibration” (Fiehn, 2006, 2008).

Analysis of metabolites using liquid chromatography/mass spectrometry (LC-MS) Metabolites were extracted from 6 mg of fresh weight ground sample using a protocol adapted from the literature (Kim *et al*., 2008). Briefly, 1.6ml of a mix of Methanol/H2O/Acetone/TFA (40/32/28/0.05, v:v:v:v) and 300ng of Apigenin (used as internal standard) were added to each sample, which was then stirred at 4°C for 30 minutes. After centrifugation (10 min, 20,000 g, 4°C), the supernatant was collected, and the pellet was extracted again by stirring with 1.6ml of the previous solvent mix for 30 minutes. After centrifugation the two supernatants were pooled, dried and resuspended in 200μl of water (ULC/MS grade)/Acetonitrile (90/10) (Biosolve Chimie, Dieuze, France) and filtered (filter paper grade GF/A Whatman®).

Metabolomics data were acquired using a UHPLC system (Ultimate 3000, Thermo Scientific, Waltham, USA) coupled to quadrupole time of flight mass spectrometer (Q-Tof Impact II Bruker Daltonics, Bremen, Germany). A Nucleoshell RP18 plus reversed-phase column (2 x 100 mm, 2.7 μm; Macherey-Nagel, Hoerdt, France) was used for chromatographic separation. The mobile phases used for the chromatographic separation were (A) 0.1% formic acid in H_2_O and (B) 0.1% formic acid in acetonitrile. The flow rate was of 400μl min^-1^ and the following gradient was used: 95% of A for 1-min, followed by a linear gradient from 95% A to 80% A from 1 to 3-min, then a linear gradient from 80% A to 75% A from 3 to 8-min, a linear gradient from 75% A to 40% A from 8 to 20-min. 0% of A was hold until 24-min, followed by a linear gradient from 0% A to 95% A from 24 to 27-min. Finally, the column was washed by 30% A for 3.5min then re-equilibrated for 3.5-min (35-min total run time). Data-dependent acquisition (DDA) methods were used for mass spectrometer data in positive and negative ESI modes using the following parameters: capillary voltage, 4.5kV; nebuliser gas flow, 2.1 bar; dry gas flow, 6L min^-1^; drying gas in the heated electrospray source temperature, 140°C. Samples were analysed at 8Hz with a mass range of 100 to 1500 m/z. Stepping acquisition parameters were created to improve the fragmentation profile with a collision RF from 200 to 700 Vpp, a transfer time from 20 to 70 µs and collision energy from 20 to 40eV. Each cycle included an MS full-scan and 5 MS/MS CID on the 5 main ions of the previous MS spectrum.

### Soluble sugars and starch analysis

Soluble sugars (sucrose, glucose, and fructose) were extracted from 150 mg of frozen tissue powder (both roots and leaves separately) with ethanol and water solution (800μl at 80% of ethanol) by incubation in a water bath at 80°C for 15 min, with shaking every 5-minute. The sample was centrifuged (5 min, 5,000 rpm) and the supernatant collected. The extraction was repeated using the same conditions with 800μl of a 50% ethanol solution, 800μl of 100% water and then 800μl 80% ethanol solution. All supernatants were mixed and then evaporated. The dried sample was resuspended in 1ml of water and kept in the dark at -20°C until soluble sugar analysis.

The residual pellet from sugar extraction was used for starch analysis. Immediately after removal of the last supernatant, the pellet was resuspended in 3ml of thermostable α-amylase. The tube was plunged in a boiling water bath and mixed every 2 mins. After 6 min, the sample was transferred to 50°C bath and 0.1ml of amyloglucosidase (20U) was added, the tube was mixed and incubated for 30 min. The entire content of the tube was then transferred to a larger tube and volume adjusted to 20ml with distilled water, mixed thoroughly, and centrifuged at 5,000 rpm for 5 min. The clear, undiluted supernatant was used for the determination of glucose issued from starch hydrolysis.

Sugar analyses were performed using Sucrose, D-Fructose and D-Glucose assay kit (Megazyme K-SUFRG; Neogene, Mi, USA) and Total starch HK assay kit (Megazyme K-TSHK), respectively, as described by the provider. Spectrophotometric measurements were conducted at 340nm in cuvettes with a 1 cm optical path.

### Gene Expression Analysis

For RNA extraction, plant material was ground in liquid nitrogen. Total RNAs from 100mg was then isolated using RNAzol® RT (Sigma), quantified, and analysed on nanodrop and 1.5% agarose gel electrophoresis to assess the purity. DNA digestion (RQ1 RNAse-free DNAse) and reverse transcription (GoScript™ Reverse Transcription) were performed as described by the manufacturer (Promega, Madison, USA). qPCR was performed (qPCR Master Mix plus CXR; Promega) using cDNA template and each set of primers. *Pathogenesis Related 1* (*PR1*; MtrunA17_Chr2g0295371) was used as SA defence gene marker and the *Proteinase Inhibitor* (*PI*; PSI-1.2; MtrunA17_Chr4g0014461) as JA pathway activation gene (Pandharikar *et al*., 2020). Other genes of interest analysed were *Chalcone Isomerase* (*CHI*; MtrunA17_Chr1g0213011), *Flavanol Synthase/Flavanone 3-Hydroxylase* (*FLS/F3H*; MtrunA17_Chr3g0092531), *Isoflavone 4’-O-methyltransferase* (*HI4’O-MT*; MtrunA17_Chr4g0046341), *D-pinitol Dehydrogenase* (*OEPB*; MtrunA17_Chr6g0480011), *Phenylalanine Ammonia-lyase* (*PAL*; MtrunA17_Chr1g0181091), *Pterocarpan Synthase 1* (*PTS*; MtrunA17_Chr7g0259091) and *SAR Deficient 4* (*SARD4*; MtrunA17_Chr1g0202471). Real-time qPCR was performed with specific primers (Table S1) using 95°C for 3 min followed by 40 cycles at 95°C for 3 sec and 60°C for 30 sec, and melting curves from 65°C to 95°C in increments of 0.5°C (AriaMx Real-time PCR machine, Agilent). Cycle threshold values (Ct) were normalized to the average Ct of two housekeeping genes *MtC27* (MtrunA17_Chr2g0295871) and *a38* (MtrunA17_Chr4g0061551) genes (Del Giudice *et al*., 2011). The expression of these two genes was not affected by the treatments. The original Ct obtained (Ariamix software; Agilent) (Dataset S1) were further used in the R qPCRBASE package (Hilliou, 2013). For each gene, the expression level of the aphid infested plants was compared with that of the non-infested control plants. The results of the qPCR analysis were generated from four independent biological repeats.

### Metabolomics Data Processing

For GC-MS, raw datafiles were converted in NetCDF format and analysed with AMDIS software (http://chemdata.nist.gov/mass-spc/amdis/). A home retention indices/mass spectra library built from the NIST, Golm (http://gmd.mpimp-golm.mpg.de/) and Fiehn databases (https://fiehnlab.ucdavis.edu), and standard compounds was used for metabolites identification. Peak areas were also determined with the Targetlynx software (Waters) after conversion of the NetCDF file in masslynx format. AMDIS, Target Lynx in split-less and split 30 modes were compiled in one single Excel file for comparison. After blank mean subtraction peak areas were normalized to Ribitol and leaves fresh weight.

For LC-MS, data files were converted and treated as previously described (Boutet *et al*., 2022). A first search was done in library with open-source software MZmine2 (Pluskal *et al*., 2010) with identification module and “custom database search” to begin the annotation with our library, currently containing 159 annotations (RT and *m/z*) in positive mode and 61 in negative mode, with RT tolerance of 0.3 min and m/z tolerance of 0.0025 Da or 6 ppm. Molecular networks were generated with MetGem software (Olivon *et al*., 2018) (https://metgem.github.io) using the .mgf and .csv files obtained with MZmine2 analysis. ESI- and ESI+ molecular networks were generated using cosine score thresholds of 0.8 in two modes. Metabolites annotation was performed as described (Boutet *et al*., 2022).

### Statistical Analysis

Statistical analysis was performed with TMEV (https://sourceforge.net/projects/mev-tm4/) for GC-MS. Univariate analyses by permutation (1-way-anova and 2-way anova) were firstly used to select the significant metabolites (P-value < 0.01). Multivariate analyses (hierarchical clustering an principal component analysis) were then made on both LC-MS and GC-MS data. Mapman (http://www.gabipd.org/projects/MapMan/) was used for graphical representation of the metabolic changes after Log2 transformation of the mean of the 3 replicates (Fiehn, 2006, 2008).

Tukey’s multiple comparison tests were also performed on independent metabolites to identify statistical differences between the various conditions for both primary and secondary metabolites. Statistical analysis of individual metabolites was done using Prism v9.1.1 (GraphPad software, USA). All experimental data are expressed as mean ± standard error (SE). Venn diagram was created based on results of Tukey’s tests, clustering metabolites based on statistical significance (P-value ≤ 0.05) using InteractiVenn (Heberle *et al*., 2015).

## Results

### Aphid infestation, more than nitrogen source, significantly alters leaf metabolite profile

*M. trunculata* leaves metabolites were analysed in four experimental conditions: nitrate-fed plants (NI), nitrate-fed plants infested with aphids (NI_Amp), nitrogen-fixing symbiotic plants (NFS) and nitrogen-fixing symbiotic plants infested with aphids (NFS_Amp) (Fig. S2). The leave extracts were analysed by GC-MS, to obtain mainly primary metabolites, and by LC-MS, to have access to the secondary metabolites. After GC-MS, 237 compounds were retained and 126 could be identified with confidence (Dataset S2), and among them the five main quantitative compounds were sucrose, phosphate, malate, citrate, and glutamate (Fig. S3). By LC-MS, 2627 compounds (in positive and negative modes) were obtained and 213 could be identified (Dataset S2). We observed a strong effect of aphid infestation and nitrogen source in the accumulation of metabolites and the hierarchical clustering of these compounds separates both nitrogen source and plant infestation status (see Fig. 1A, S4 and S5 for GC-MS and Fig. S6 for LC-MS, respectively). From the four experimental conditions, a total of 194 unique metabolites, 126 from GC-MS and 69 from the LC-MS, were found significantly different according in their accumulation (Dataset S2). Only sucrose was found significantly in both analysis. Among the 194 identified the most represented classes were flavonoids with 40 different compounds, followed by 30 carboxylic acids, 26 amino acids and 19 sugars (Fig. 1b).

**Fig. 1:**
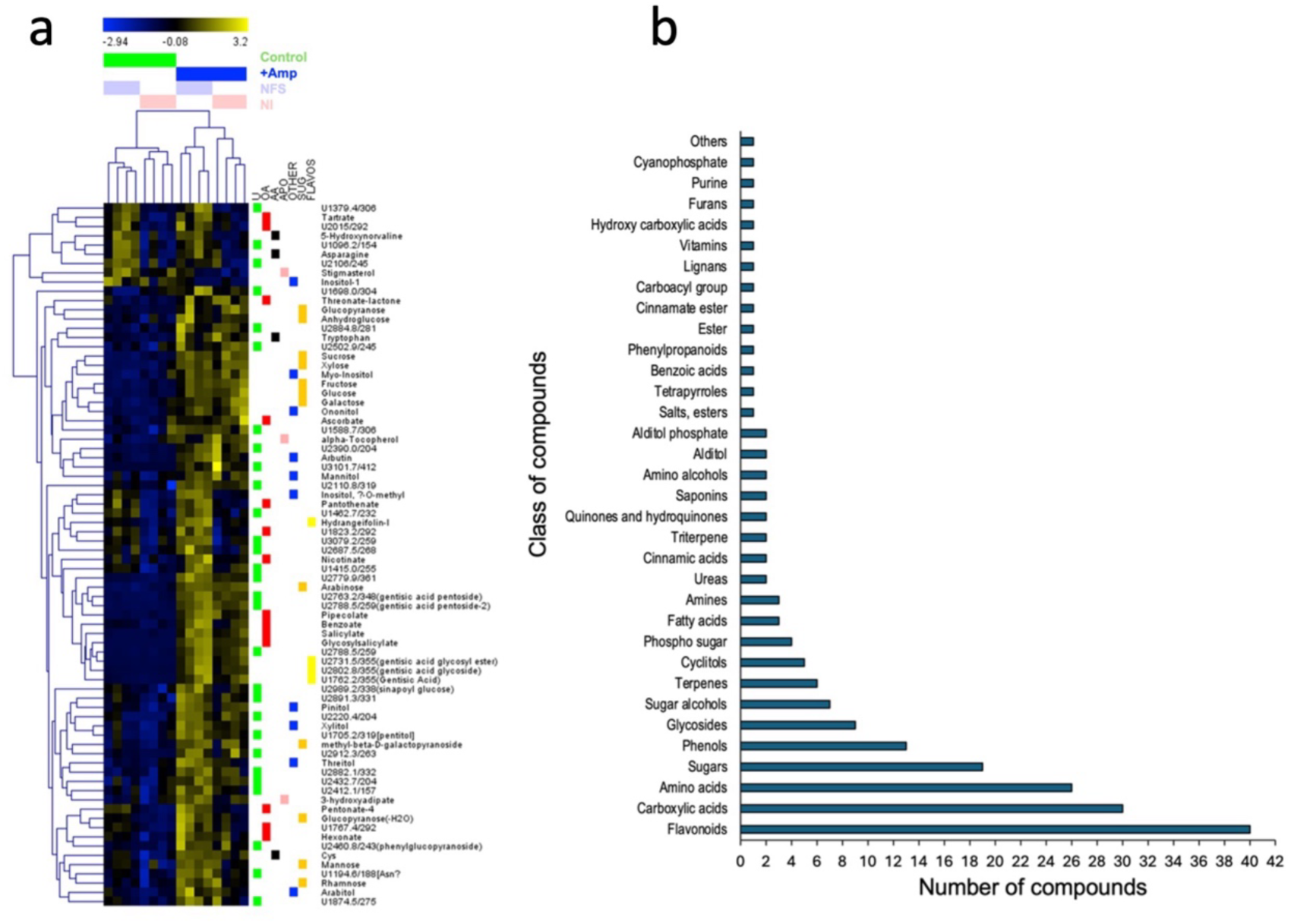
Aphid infestation, more than nitrogen source, significantly alters leaf metabolite profile. a) Heatmap from GC-MS analysis showing unsupervised hierarchical clustering of compounds separated by both infestation and nitrogen source. b) Class abundance of metabolites from both GC-MS and LC-MS analyses.

We then identified statistical differences for the 194 unique metabolites between the four experimental conditions using multiple comparison tests (Fig 2; Dataset S3). 86 compounds were not found significatively accumulated in one specific condition, including many of the amino acids. 108 compounds were found to be differential accumulated in the different conditions. Amongst them, aphid infestation increased significantly the accumulation of 62 metabolites. 18 metabolites accumulated in control NFS conditions and in infected conditions, suggesting they are linked to a general infection/infestation response. 2 compounds are specifically found in NFS conditions and 19 compounds were significantly accumulated in NFS_Amp conditions.

**Fig. 2:**
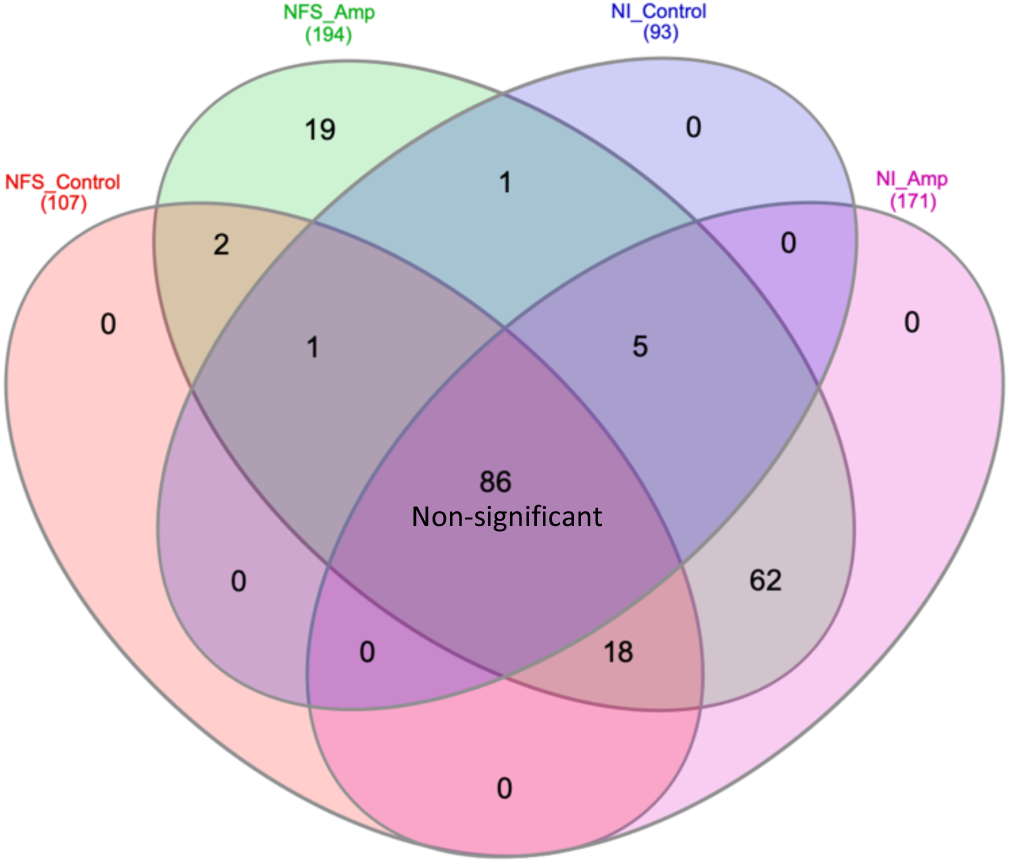
Metabolite distribution according to nitrogen source and aphid infestation affects the metabolite profile of plants. Venn diagram showing distribution of statistically significant metabolites across the various experimental conditions. The name of the compounds and the statistic can be found in Dataset S3.

### Aphid infestation triggers a greater accumulation of sugars than amino acids in both NFS and NI Plants

#### Amino Acids

The majority of the amino acids (20) were not significantly accumulated in any specific condition. In contrast, tryptophan, cysteine, asparagine were significantly accumulated in NFS control, NFS and NI Amp. 5-Hydroxynorvaline was significantly accumulated in NFS control and NFS Amp compared to NI control and NI Amp. Finally, the putative asparagine or asparagine derivative (U1194.6/188[Asn?) is significantly more accumulated in NFS and NI Amp than in NFS control and NI control (Dataset S3).

#### Sugars

Among the 62 metabolites significantly accumulated under the aphid-infested conditions, glucose, sucrose and fructose contents were increased between three to five-fold compared to control conditions in leaves (Fig. 3). To investigate whether this leaf increase in sugar was associated with either a reduction in sugar transport from leaf to root or the results of the mobilization of leaf starch, sugar and starch contents were measured using biochemical assays in the leaves and roots of plants under the different conditions (Fig. 4). The increase in glucose, sucrose and fructose in infested leaves was confirmed (Fig. 4), but no significant accumulation of starch in leaves and no change in root sugar concentration were observed, whatever the plant’s growth conditions (Fig. 4), suggesting an alteration of the leaf sugar metabolism in response to aphid infestation.

**Fig. 3:**
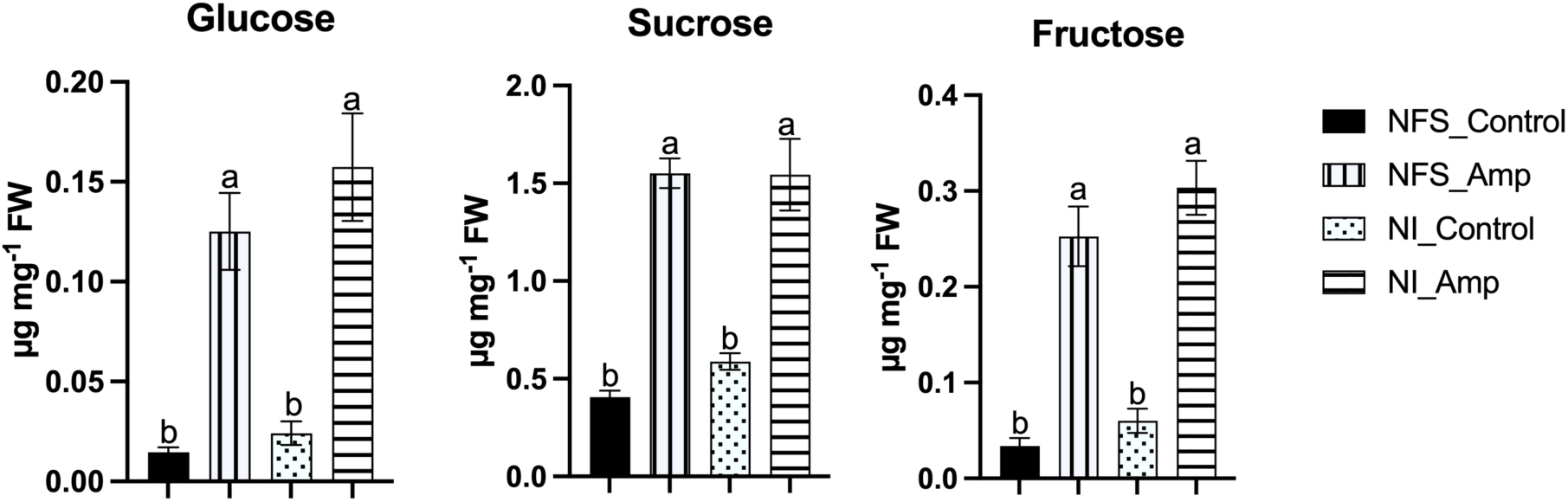
Aphid infestation triggers accumulation of soluble sugars in both NFS and NI plant leaves. Bar graphs from GCMS analysis showing the significant accumulation of glucose sucrose and fructose in NFS and NI plants upon aphid infestation. Metabolites analysed and quantified in μg mg^-1^ of fresh weight (FW) of leaves of NFS and NI plants 12 days after infestation by the aphids (control = no aphid infestation). Mean ± SE (n=4): different letters indicate a *p* ≤ 0.05 (Tukey’s test).

**Fig. 4:**
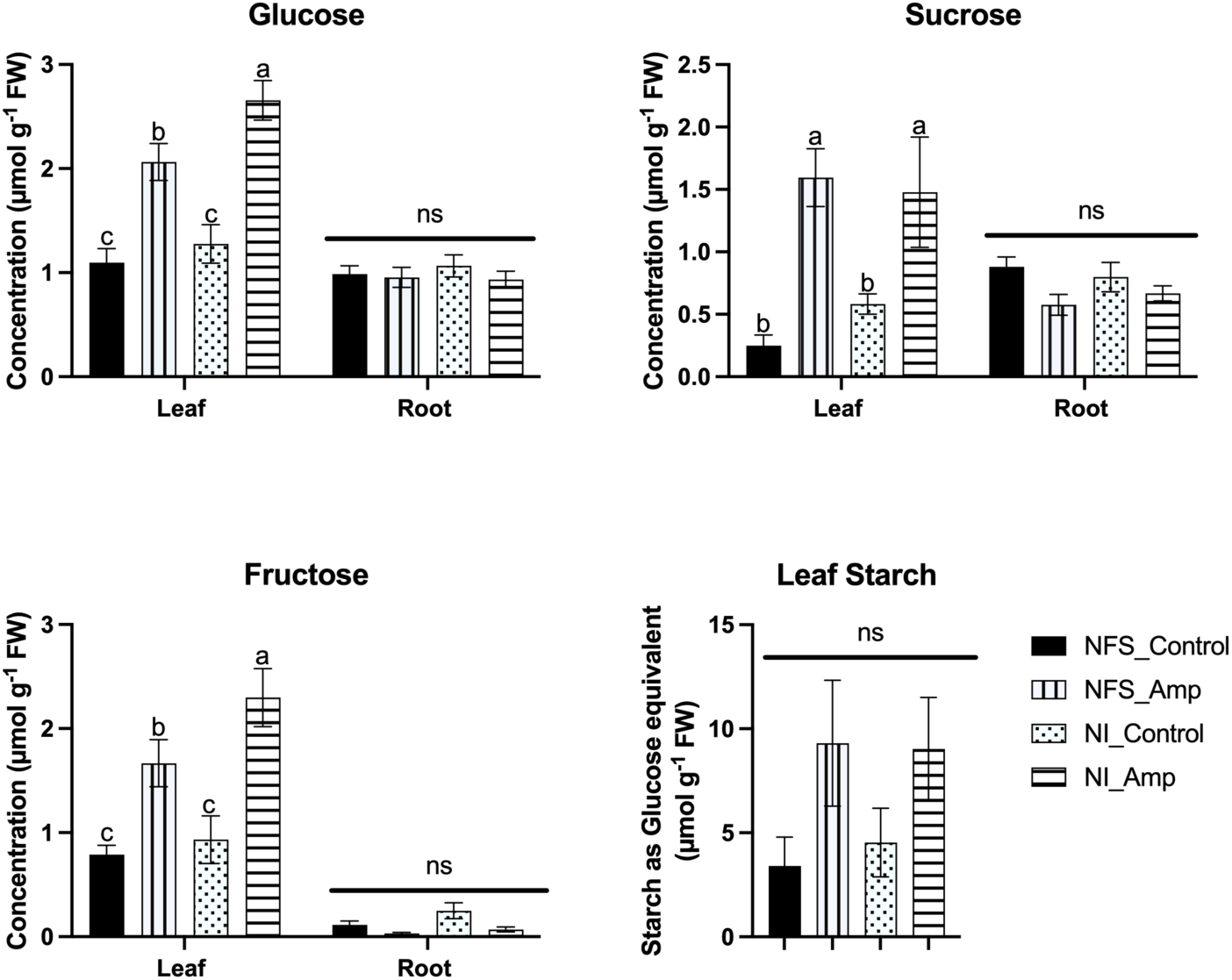
Aphid infestation affects sugar concentration in leaves but not in roots. Biochemical analysis showing the significant accumulation of glucose, sucrose, and fructose in leaves of NFS and NI plants. No significant difference was found in roots upon aphid infestation. No statistical difference was observed in leaves starch, measured as glucose equivalent, between the different conditions. Metabolites analysed and quantified in μmol g^-1^ of fresh weight (FW) of leaves and roots from NFS and NI plants 12 days after infestation by the aphids (control indicates no aphids infestation). Mean± SE (n=4); different letters indicate a *p* ≤ 0.05 (Tukey’s test).

### Aphid infestation triggers accumulation of defence related metabolites in both NFS and NI plants

Amongst the metabolites significantly accumulated during aphid infestation (Fig. S5; Dataset S3), many are involved in the regulation of plant defence pathways such as salicylates, pipecolate, an intermediate of the lysine catabolic pathway, pinitol, a cyclitol derived from myo-inositol, as well as secondary metabolites with known defence activity such as the putative daidzin, the glycoside form of the aglycone daidzein (Fig. 5).

**Fig. 5:**
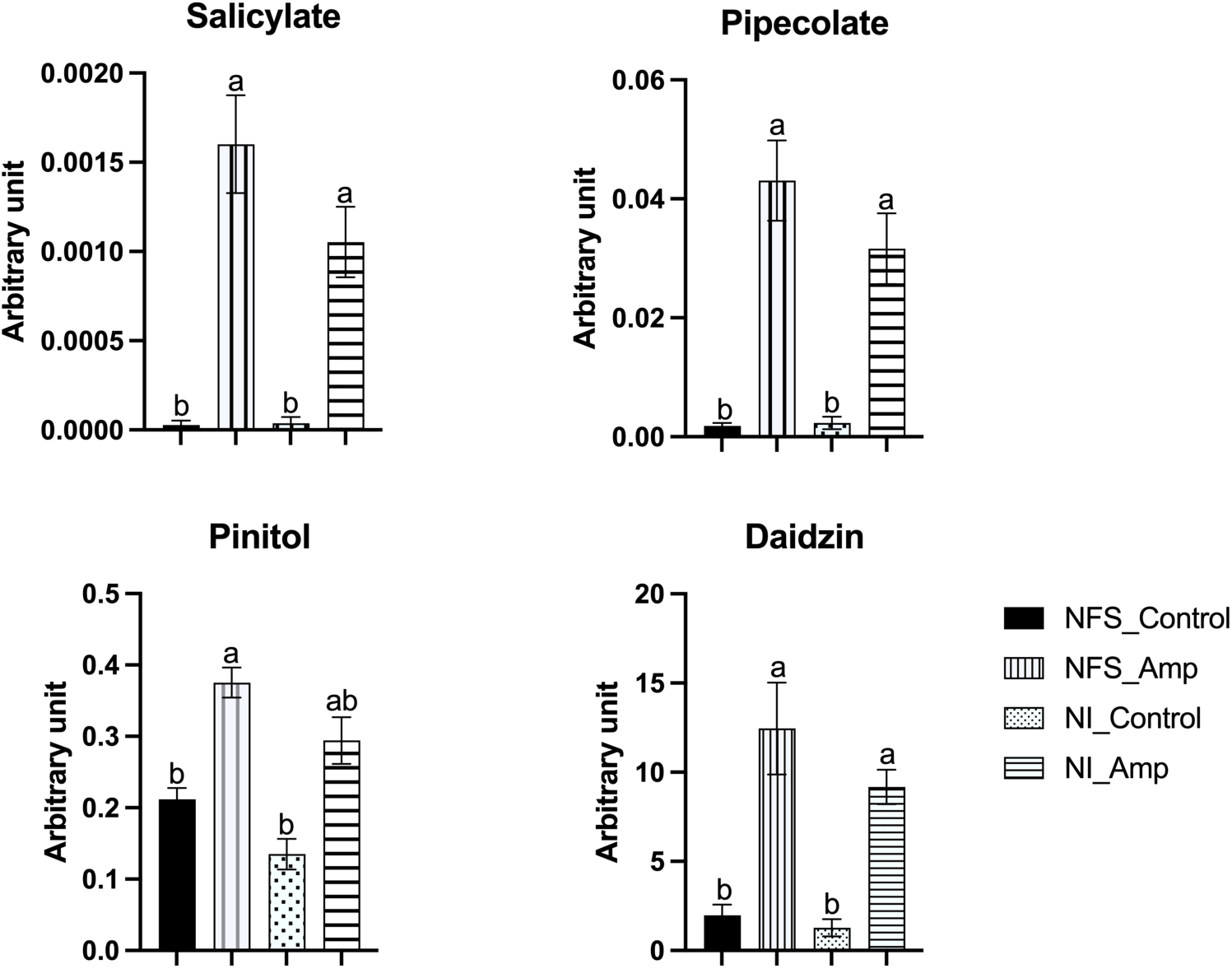
Aphid infestation triggers accumulation of defence-related metabolites in both NFS and NI particularly from the salicylic acid defence pathway. Graphs show accumulation of metabolites expressed as arbitrary unit (in mg apigenin equivalent mg^-1^) of leaves fresh weight (FW) of NFS and NI plants 12 days after infestation by the aphids (control indicates no aphids infestation). Mean± SE (n=4); different letters indicate a *p* ≤ 0.05 (Tukey’s test).

### Aphid infestation induces expression of genes involved in the flavonoid synthesis pathway

In order to test whether pipecolate, pinitol, and diadzin accumulation resulted also from an increase in expression of the gene(s) involved in their synthesis pathways, we have measured the expression of *Chalcone isomerase (CHI)*, *flavonol synthase/ flavonone 3β-hydroxylase (FLS/F3H)*, *hydroxyisoflavone-O-methyl transferase* (*HI4’O-MT)* and *Pterocarpan synthase* (*PTS)*, *Phenylalanine Ammonia Lyase (PAL), SAR-DEFICIENT4 (SARD4)* and (*OEPB*) by RT-qPCR (Table 1). Since we previously showed that *PR1,* a marker for SA defence pathway and *PI*, a marker for jasmonic acid defence pathway were also differently induced in NFS and NI plants after aphid infestation (Pandharikar *et al*., 2020), we test these genes as the plant condition controls. Here, RT-qPCR analysis showed that *PR1* expression increased 3.4 times more in NI_Amp condition than in NFS_Amp condition. In contrast *PI* was 2.3 times more expressed in NFS_Amp condition than in NI_Amp condition, in agreement with our previous results. The analysis of the expression of genes involved in secondary metabolism pathway showed that no significant change in *SARD4* and *PAL* expression was detected in aphid infected compared to the control plants. *OEPB* expression was significantly induced in NI_Amp leaves but not NSF_Amp leaves, although the amplitude of increase was similar. *CHI*, *FLS/F3H*, *HI4’O-MT* and *PTS* showed significant induction upon aphid infestation in both NFS and NI conditions; the induction being 2-fold for *FLS/F3H*, and reaching more than 50-fold for *PTS* in NSF_Amp.

**Table 1:** Gene expression analysis of NFS and NI plants. Rescaled qPCR expression showing the level of gene induction upon aphid infestation in NFS and NI plants (NFS_Amp and NI_Amp, respectively) compared to their control (NFS and NI, respectively). Data are expressed as mean ± standard error (SE); t-test on all genes, P ≤ 0.05, significant.

| Gene | Pathway | Product | Fold Increase from Control<br>(Mean ± SE) | | *p* value | |
| --- | --- | --- | --- | --- | --- | --- |
|  |  |  | NFS_Amp/<br>NFS | NI_Amp/NI | NFS_Amp/<br>NFS | NI_Amp/NI |
| *PR1* | SA pathway | Salicylic acid | 3.5 ± 0.32 | 11.99 ± 1.25 | 0.05 | 0.02 |
| *PI* | JA pathway | Jasmonic acid | 10.72 ± 0.26 | 4.73 ± 0.75 | 0.0006 | 0.002 |
| *PAL* | Phenylpropanoid pathway | Phenylpropanoids | 1.40 ± 0.02 | 1.35 ± 0.14 | 0.06 | 0.31 |
| *SARD4* | Lysine Biosynthesis pathway | Pipecolate | 1.16 ± 0.29 | 0.74 ± 0.09 | 0.73 | 0.88 |
| *OEPB* | Pinitol Biosynthesis | Pinitol | 1.75 ± 0.01 | 1.96 ± 0.31 | 0.07 | 0.01 |
| *FLS/F3H* | Isoflavonoid pathway | Flavonoids | 1.96 ± 0.05 | 2.5 ± 0.40 | 0.00002 | 0.01 |
| *HI4'O-MT* | Isoflavonoid pathway | Formononetin | 4.32 ± 0.10 | 3.37 ± 0.53 | 0.001 | 0.02 |
| *CHI* | Isoflavonoid pathway | Flavonoids | 4.58 ± 0.11 | 5.83 ± 0.93 | 0.006 | 0.004 |
| *PTS* | Isoflavonoid pathway | Medicarpin | 53.35 ± 1.27 | 24.66 ± 3.91 | 0.0001 | 0.001 |

Rescaled qPCR expression showing the level of gene induction upon aphid infestation in NFS and NI plants (NFS\_Amp and NI\_Amp, respectively) compared to their control (NFS and NI, respectively). Data are expressed as mean ± standard error (SE); t-test on all genes, $P \leq 0.05$ , significant.

Taken together these results showed that the accumulation of secondary metabolites is at least partially associated to an higher expression of enzymes involved in their synthesis pathways.

### NFS induces a differential defence mechanism upon aphid infestation

Nineteen metabolites (10% of the identified metabolites) were significantly accumulated in NSF_Amp plants compared to other conditions (Fig. S7; Dataset S3). Fifteen of them are secondary metabolites from phenyl-propanoid (13) and terpene (2) synthesis pathways. For example, amongst these metabolites, putative triterpenoid saponin 3-Glu(1-3)Glu-28-Xyl(1-4)Rha(1-2)Ara zanhic acid is increased two-fold in NFS_Amp plants compared to NI-Amp plants, putative 3-(4’O-Malonyl)Rha(1-2)Gal(1-2)GluA-Soyasaponenol B is 2.8 time more accumulated in infested NFS-Amp plants than in the other three plant growth conditions, flavonoid putative tricin 5-glucoside is about 2 times more accumulated in NFS-Amp compared to NI-Amp while glycosylsalicylate is half a fold more in NFS-Amp than in NI-Amp (Fig. 6).

**Fig. 6:**
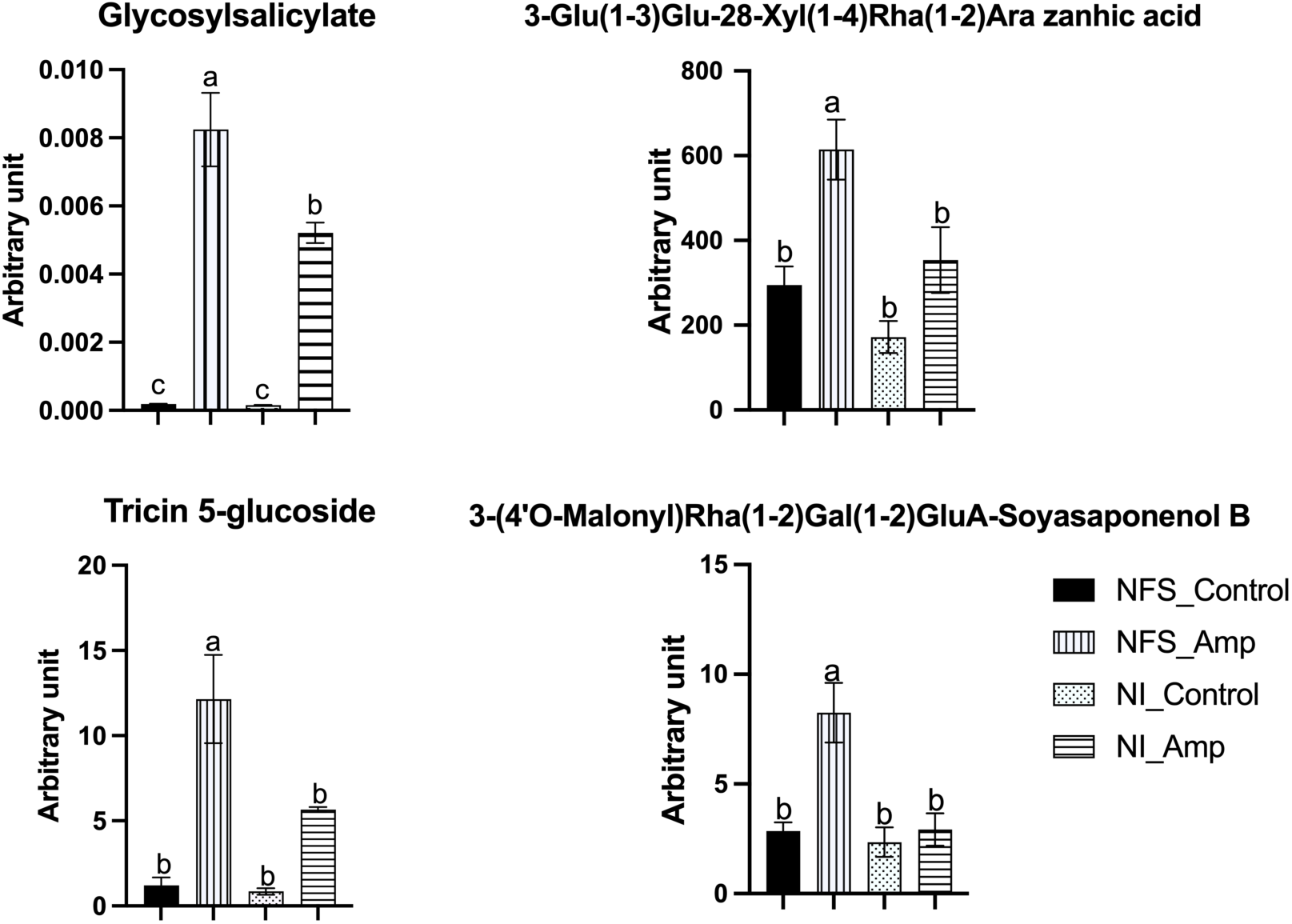
NFS induces a differential defence mechanism upon aphid infestation. Graph showing significantly accumulated metabolites in NFS upon aphid infestation. Metabolites expressed as arbitrary unit (in mg apigenin equivalent mg^-1^) of leaves fresh weight (FW) of NFS and NI plants 12 days after infestation by aphids (control indicates no aphids infestation). Mean± SE (n=4); different letters indicate a *p* ≤ 0.05 (Tukey’s test).

## Discussion

There is a growing interest in better understanding the roles of symbiotic microbes in plant defence and in a broader sense how this could influence plant interactions with bioagressors. Beneficial microbes, in addition to enhancing plant growth and development have been reported to induce defence reaction and confer protection on their host plants (Gopalakrishnan *et al*., 2015; Liu *et al*., 2020; Benjamin *et al*., 2022). Rhizobia are able to induce systemic resistance in legumes such as pigeon pea against fusarial wilt as it was found that a combination of rhizobial strains was better in inducing resistance (Dutta *et al*., 2008).

Upon pea infection with the fungi Didymella, Turetschek and colleagues (2017) observed a strong increase in sugars, sugar alcohols and glycolysis/TCA intermediates when studying cultivars in symbiosis with rhizobia and mycorrhiza. They also observed the accumulation of galactose, raffinose, maltose, threitol, melibiose, fructose and pyruvate in the pea Protecta cultivar. Similar accumulation was also reported for amino acid pools, however, there was significant depletion of phenylalanine in the pea cultivar Messire (Turetschek *et al*., 2017).

Plants produce diverse primary and secondary metabolites that are involved in various functions including development and defence. Previous work have demonstrated that aphids are able to modify the overall metabolite profile of plants to establish feeding and that plants may react by producing metabolites that have antifeeding or deterrent effects (Giordanengo *et al*., 2010; Kumar, 2017; Jakobs *et al*., 2019; Shih *et al*., 2023). Few metabolomic studies have focused on pea aphids and legumes interaction (Sanchez-Arcos *et al*., 2019) and none have addressed the role of rhizobia bacteria on the plant metabolomic response during aphid attack. Based on our previous observation that nitrogen-fixing symbiosis is detrimental to aphid fitness in *M. truncatula* in association with potential changes in hormonal balance between SA and JA (Pandharikar *et al*., 2020), we performed an untargeted metabolomic analysis in *M. truncatula* leaves 12 days after aphid infestation (chronic effect) in both NFS and NI plants.

In our study, among amino acids (Dataset S3), we observed an increase in asparagine content in NFS plant leaves compared to NI control plants that could be associated with asparagine formation in root nodule which account for 60% of nodule amino acid content (Sulieman & Schulze, 2010). Similarly, cysteine was found to be more accumulated in symbiotic *Lotus japonicus* plants as the nodule is an important source of reduced sulphur in the plant (Kalloniati *et al*., 2015). In contrast, tryptophan accumulation in the leaves of NFS plants has not yet been observed, and a potential role in defence mechanism can be assumed (Hiruma *et al*., 2013). In NFS- and NI-infested plants, the increase of asparagine, cysteine and tryptophan content compared to NI plants could be associated with the change of metabolism induced by aphids for better nutritional content (Chiozza *et al*., 2010; Tegeder, 2014) or by the plant through the induction of defence reaction (Taylor & Ostrowsky, 2019). Citrate, fumarate, malate and succinate accumulation were not significantly different among our four growth conditions, suggesting that tricarboxylic acid (TCA) cycle is not changed by NFS or aphid infestation. In contrast to the large number of metabolites that did accumulate differentially in the different growth conditions, two (5-hydroxynorvaline and tartrate) were specifically accumulated in NFS conditions. Whereas the significance of tartrate accumulation, which is and end point product of the catabolism of ascorbic acid, is more difficult to analyse (Burbidge *et al*., 2021), unless it is used by the rhizobial bacteria (Ramachandran *et al*., 2011), that of the 5-hydroxynorvaline, a non-protein amino acid, may be related to its defensive functions against insects (Huang *et al*., 2011; Yan *et al*., 2015). No metabolite was specifically more accumulated in NI- and infested NI-conditions.

Our data show that the sugar metabolism is particularly affected by the aphid infestation (Fig. 3). Amongst the primary metabolites, 12 are sugars and represent 70% of the number of primary metabolites significantly accumulated in infested plants (Dataset S3). In our experimental conditions, we were interested by the sugar transport to root to feed the root nodule involved in biological nitrogen fixation. Indeed, we previously showed that biological nitrogen fixation is affected in the infested conditions (Pandharikar *et al*., 2020) and we hypothesized that the decrease in biological nitrogen fixation could be associated to a reduced sugar transport from leaves to roots. However we did not observed a significant modification in the accumulation of root glucose, fructose and sucrose between control and infested plants, indicating that the modification of sugar metabolism is not associated with the sugar transport to the root. We also observed that starch seemed more accumulated in infested leaves than in control ones but this difference was not significant. In conclusion, the modification of sugar metabolism does not seem to be associated to a modification of sugar storage through starch or sugar transport to roots. Moreover, the modification of sugar metabolism by aphids seems not dependent of the nitrogen source. The accumulation of sugars has been previously reported in plant-aphid (Ponzio *et al*., 2017) as well as in plant-pathogen interactions (Kanwar & Jha, 2019) as being involved in the coordination of plant defence signalling (Yamada & Mine, 2024), this perhaps could be an explanation for our observation.

A large number of significantly accumulated metabolites (62 metabolites, 31% of the total metabolites) is present in NFS- and NI-infested plants compared to their control counterparts. This results show that Medicago plants respond significantly to aphid infestation and that the nitrogen source does not play a significant role in their accumulation. Amongst these secondary metabolites, 72% are from 3 families, flavonoids (30), phenolics (11) and glycosides (4). These different compounds are mainly associated with plant defence against pests such as biocide activity (i.e. acacetin, chrysoeriol, daidzin), feeding deterring activity and defence signalling activity (i.e. salicylate, pipecolate) (Stochmal *et al*., 2001; Goławska *et al*., 2012, 2024; Kim *et al*., 2022; Pawłowska & Stepczyńska, 2022). Pinitol has also been shown to participate to the biological control of powdery mildew in cucumber (Chen *et al*., 2014) and myoinositol influences the plant bacterial colonization (O’Banion *et al*., 2023). Thus, in parallel with secondary metabolism, the modification of inositol metabolism in NFS plant may be involved in the differential defence process observed in infested NFS-plants compared to compared to infested NI-plants. The significant differential accumulation of 3-Glu(1-3)Glu-28-Xyl(1-4)Rha(1-2)Ara zanhic acid and 3-(4’O-Malonyl)Rha(1-2)Gal(1-2)GluA-Soyasaponenol B, two terpene molecules are also a marker of both the symbiotic state and the infection by aphids. Finally, 3-(4’O-Malonyl)Rha(1-2)Gal(1-2)GluA-Soyasaponenol B maybe one of the most interesting and intriguing metabolite in our experiment. This triterpenoid saponin is a defensive compound against pathogenic microbes and herbivores (Osbourn, 1996; Kuzina *et al*., 2009; Szakiel *et al*., 2011) and may act as feeding deterrents for plant specialist herbivores (Cui *et al*., 2019) causing a cytotoxic effect in hindgut and fat body of insects (Adel & Sammour, 2012). Production of triterpenoids have also been observed as an effect of rhizobia on pea seeds infected with Didymella. Accumulation of seed terpenoid Pisumoside B was observed in uninfected rhizobial treated seeds, meanwhile, Soyasapogenol C, Api_Dai_Kae_Flavon and 6-Hydroxyapigenin 7-[6”-(3-hydroxy-3-methylglutaryl)glucoside] were significantly enhanced in infected rhizobial treated seeds in Protecta cultivar (Ranjbar Sistani *et al*., 2017). It may represent a very interesting biological marker of the symbiosis induced defence priming. The induction of defence mechanisms is associated to reprograming of gene expression in plants (Aerts *et al*., 2022). For example reciprocal interactions between a chewing herbivore, *Sitona lineatus* (pea leaf weevil) and *Pisum sativum* (pea) plants grown with or without rhizobia (*Rhizobium leguminosarum* biovar. *viciae)* revealed that plants grown with rhizobia had increased gene transcript expression associated with hormone-related defence (jasmonic acid, ethylene, abscisic acid) as well as physical and antioxidant-related defence, which may explain reduced feeding by *S. lineatus* (Basu *et al*., 2022).

We have also looked at the expression of some genes involved in the secondary metabolism pathway. As expected from previous analysis, *PR1* and *PI* are differentially expressed in infested NFS plants and NI plants suggesting that different defence transcriptional reprograming occurs in the two conditions.

Looking at the genes involved in secondary metabolism synthesis pathway, transcript accumulation of *SARD4*, that encodes a key enzyme for pipecolic acid biosynthesis, and *PAL*, the first enzyme of the phenylpropanoid pathway, is not modified by aphid infestation neither in NFS-plants nor in NI-plants. This was surprising since *SARD4* was shown to be required for the establishment of systemic acquired resistance to pathogen infection in Arabidopsis (Ding *et al*., 2016). However, *SARD4* deleted plants are still able to biosynthesize pipecolate, this pathway may involve thus other enzyme(s). PAL is also a member of a multigenic family and other members of this family may be induced. In contrast the isoflavonoid pathway genes *CHI, FLS/F3H, HI4’O-MT* and *PTS* expression was significantly increased in infested plants: while the induction of *CHI, FLS/F3H, HI4’O-MT* does not vary more than two fold in the two infested growth conditions, the *PTS* expression is induced 53- and 25-fold times by aphids in NFS and NI plants, respectively. *PTS* is involved in synthesis of pterocarpans that constitute the second largest group of natural isoflavonoids and play an important role as phytoalexins. In Medicago, the pterocarpans medicarpin was shown to protect the plant from the powdered mildew *Erysiphe pisi* and activates the SA pathway (Gupta *et al*., 2022). Medicarpin was also shown to be accumulated in Medicago leaves upon long term or strong attack by pea aphids (Stewart *et al*., 2016) and also in response of infection with the fungal pathogen *Phoma medicaginis* (Jasiński *et al*., 2009) suggesting some large spectrum defensive role. In contrast, medicarpin was shown to be an antagonists of nod gene expression necessary for rhizobia to form their association with the plant roots (Zuanazzi *et al*., 1998). Thus an increase in medicarpin synthesis during aphid infestation could also explain in part the effect on the root nodules previously observed (Pandharikar *et al*., 2020). *OEPB*, a gene involved in pinitol synthesis (Pupel *et al*., 2019), was significantly induced in infested-NI plants and not in infested-NFS plants. Pinitol has been shown to prolonged the pea aphid probing behaviour but did not prevent them from feeding (Campbell & Binder, 1984; Kordan *et al*., 2011). Pinitol has also been involved in maintenance of the nodule osmotic balance during development (Poole & Ledermann, 2022) and *Sinorhizobium meliloti*, may catabolize pinitol to form nodule (Kennedy-Mendez, 2018). Thus why *OEPB* increases only in NI plant remains to be understood.

In aphid infestation, signal transduction pathway is activated to induce related transcription factors that regulate the expression of downstream disease resistance genes (Gao *et al*., 2021). Taken together, our results show that, in our conditions, aphids are able to trigger accumulation of various metabolites and expression of defence genes responsible for SAR and SA defence pathway as well as modify the metabolism of primary sugars in legumes. Under infestation by pea aphids, legumes in symbiosis are able to produce a differential defence reaction by activating the JA defence pathway and production of defence metabolites such as triterpenoid saponins (soyasapogenol). Here, we have chosen one aphid line that develops well on the A17 *Medicago truncatula* cultivar used, however we know that the metabolic profiles of plants infested with their native or non-native aphid host race could vary (Sanchez-Arcos *et al*., 2019). Meanwhile in these interactions the potential change induced by the presence of the rhizobium is not questioned since the symbiotic status of the plant used is generally unknown. Thus our results provide a foundation for the development and inclusion of symbiosis in such studies and in a more applied role as a potential stimulant of plant defence against herbivores that may be a new form of biocontrol in integrated pest management.

## Supporting information

Benjamin et al._Supplementary data

## Acknowledgements

We are highly grateful to S. Tares-Amichot and L. Arthaud for help in aphids manipulation and rearing and for M. Bosseno for help in Medicago plants seedling. GB is supported by a doctoral fellowship from the department “Santé des Plantes et Environnement” of Institut National de Recherche pour l’Agriculture, l’Alimentation et l’Environnement (INRAE) and the Université Côte d’Azur (UniCA). The IJPB benefits from the support of Saclay Plant Sciences-SPS (ANR-17-EUR-0007). This work has benefited from the support of IJPB’s Plant Observatory platform PO-Chem. This work was supported by UniCA, the SPE department of INRAE and the French Government (National Research Agency, ANR) through the “Investments for the Future” programs LABEX SIGNALIFE ANR-11-LABX-0028-01 and IDEX UCAJedi ANR-15-IDEX-01.

## Author contributions

GB, PF, MP, and JLG conceived the ideas. GB, PF, MP, and JLG planned and designed the research. SB and GC performed the metabolomics analysis and analysed the data. GB and MP performed qPCR analysis. GB and RB performed sugar and starch analysis. GB analysed all other data. GB wrote the manuscript with inputs from PF, MP, and JLG. All authors contributed and agreed to the final revisions.

## Data Availability

The metabolomics data that support the findings of this study are available in MassIVE server with the reference number MSV000095254; http://doi.org/10.25345/C5CV4C35C.

Datasets S1, S2 and S3 are available on Recherche Data Gouv server at https://doi.org/10.57745/PCDG8H.

## Supplementary Data

Datasets available as: Benjamin, Goodluck, 2024, “Supplementary Data for Differential induction of Medicago truncatula defence metabolites in response to rhizobial symbiosis and pea aphid infestation”, https://doi.org/10.57745/PCDG8H, Recherche Data Gouv, V1, UNF:6:A0SbGGbxQocqa6JsmMuoMA== [fileUNF]

Dataset S1: File containing qPCR Data (qPCR CT Data.xlsx)

Dataset S2: File containing GC-MS and LC-MS metabolomics data (Metabolomics_data.xlsx)

Dataset S3: File containing Significant accumulated metabolites analysed by Anova and Venn Analysis (Significant metabolites_Venn.xlsx)

---

**Fig. S1:**
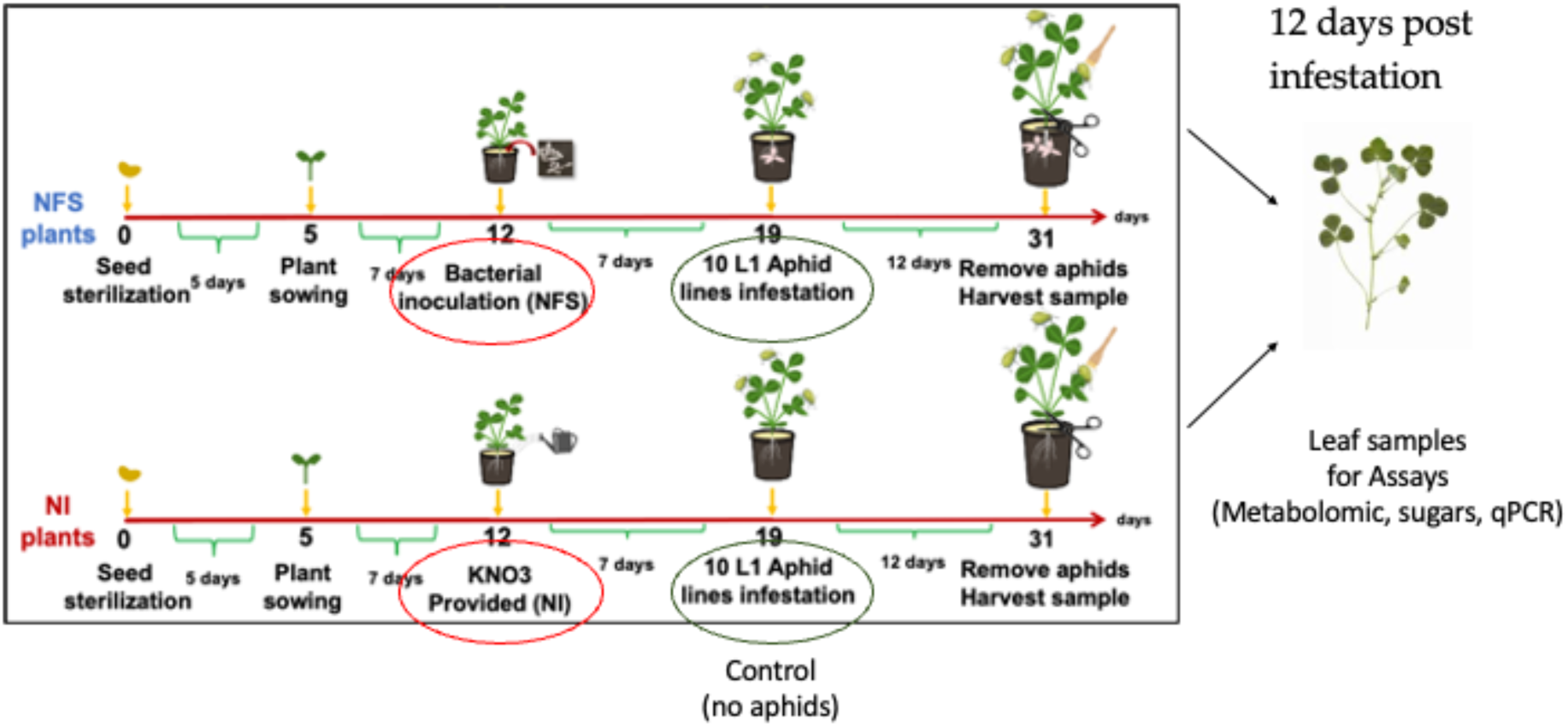
Experimental setup. Diagram showing timeline leading up to assay with key date points for bacteria inoculation and aphid infestation marked up. Each pot contained six plants and 4 pots were made for each condition per biological replicate. Assay was performed using 4 biological replicates produced using the same timeline.

**Fig. S2:**
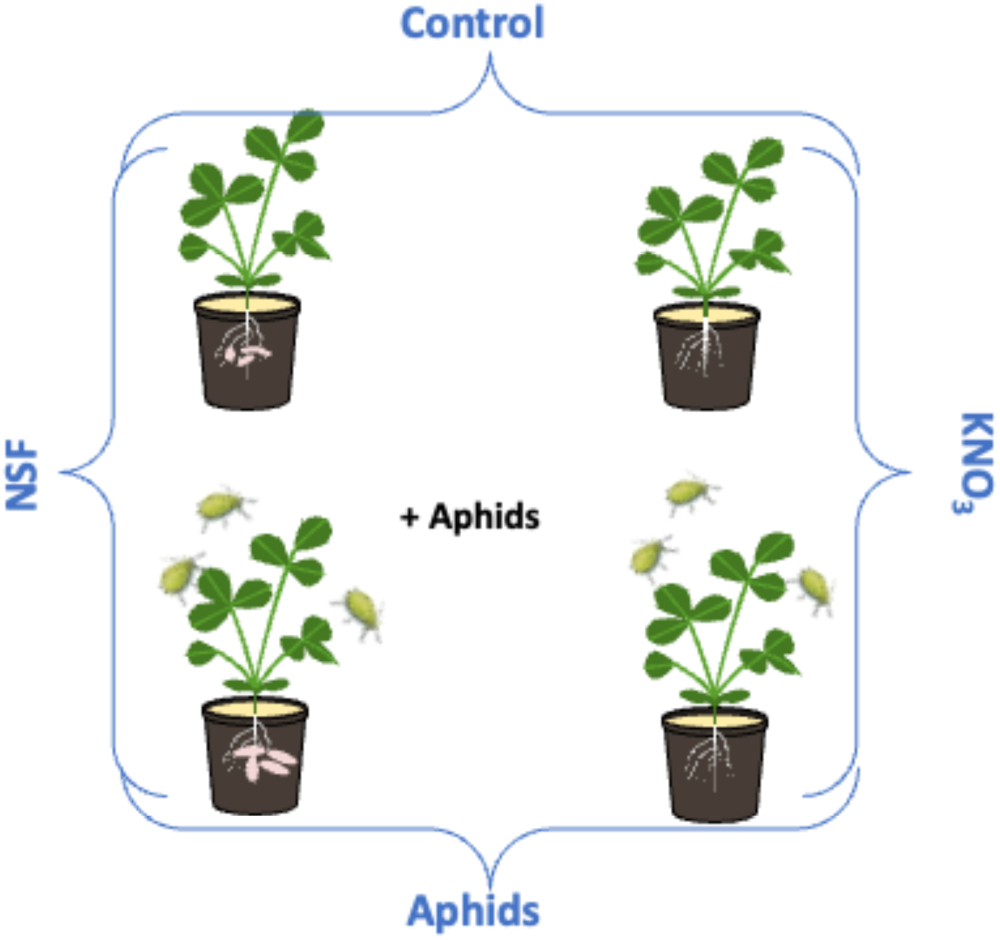
Experiment treatment classification. Diagram showing the classification of studied treatments into 4 groups, NFS_Control, NFS_Amp, NI_Control (KNO_3_) and NI_Amp (KNO_3_).

**Fig. S3:**
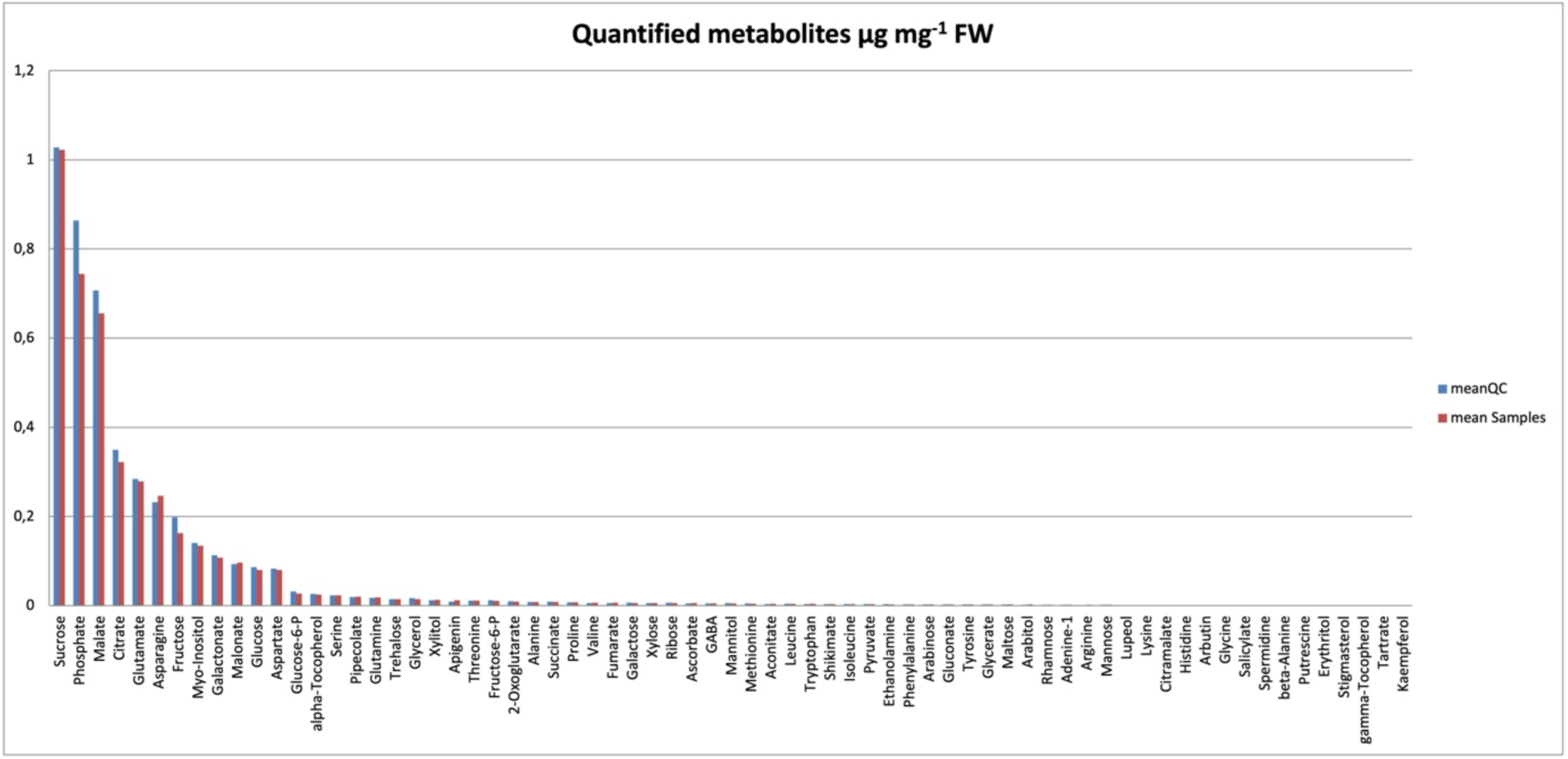
Abundance of quantified metabolites. Graph of quantified metabolites from GC-MS analysis showing accuracy of abundance by comparing the mean obtained from the quality controls (QC, blue bars) and mean obtained from the samples (Samples, red bars).

**Fig. S4:**
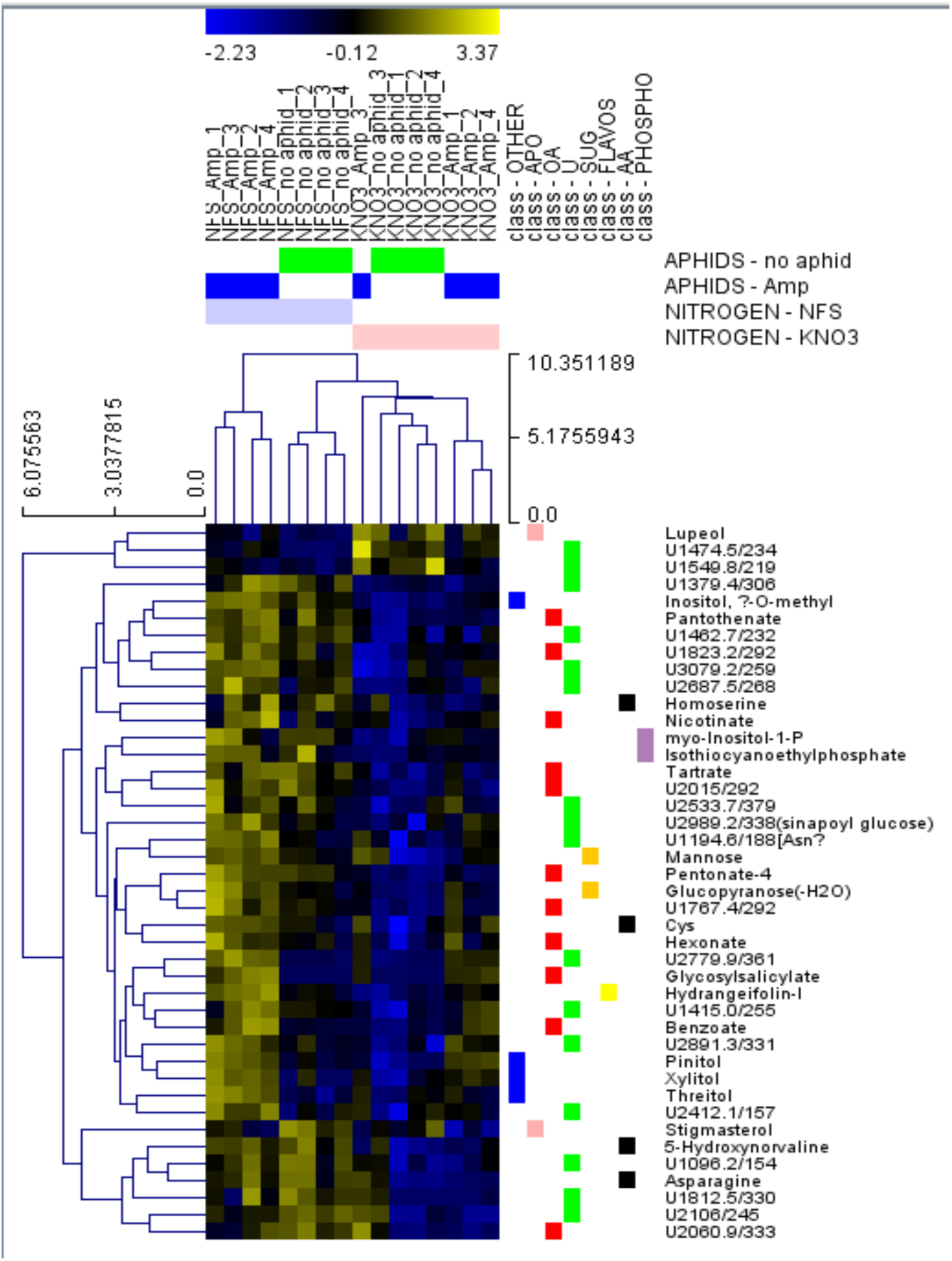
Nitrogen source significantly modifies the leaf metabolite profile. Heatmap of 2-way Anova analysis from GC-MS showing unsupervised hierarchical clustering of compounds separated by nitrogen source. NFS = Nitrogen fixing symbiosis, KNO3 = Non inoculated, Amp = aphid infested, no aphids = Control.

**Fig. S5:**
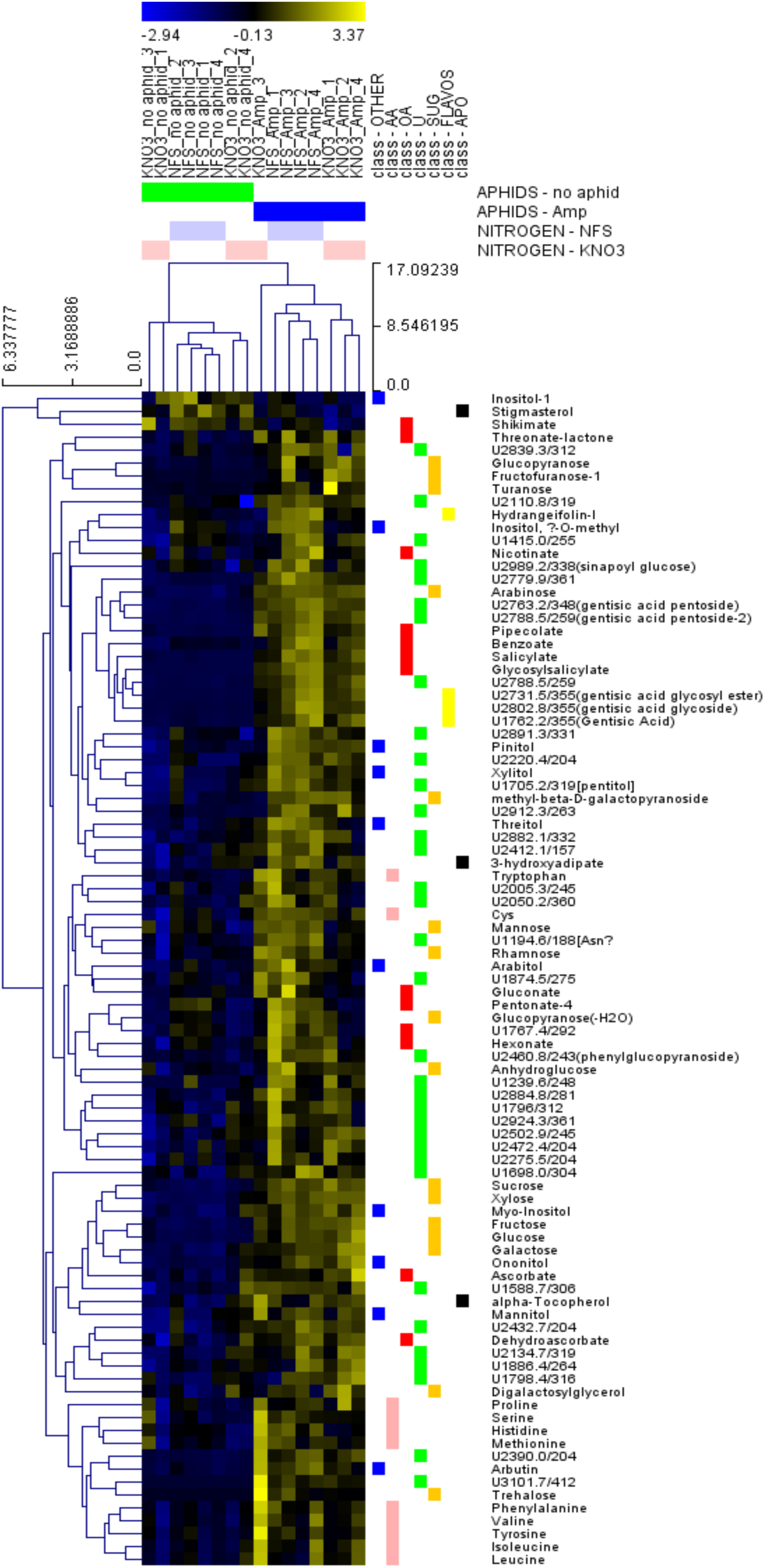
Aphid infestation significantly modifies the leaf metabolite profile. Heatmap of 2-way Anova analysis from GC-MS showing unsupervised hierarchical clustering of compounds separated by aphid infestation. NFS = Nitrogen fixing symbiosis, KNO3 = Non inoculated, Amp = aphid infested, no aphids = Control.

**Fig. S6:**
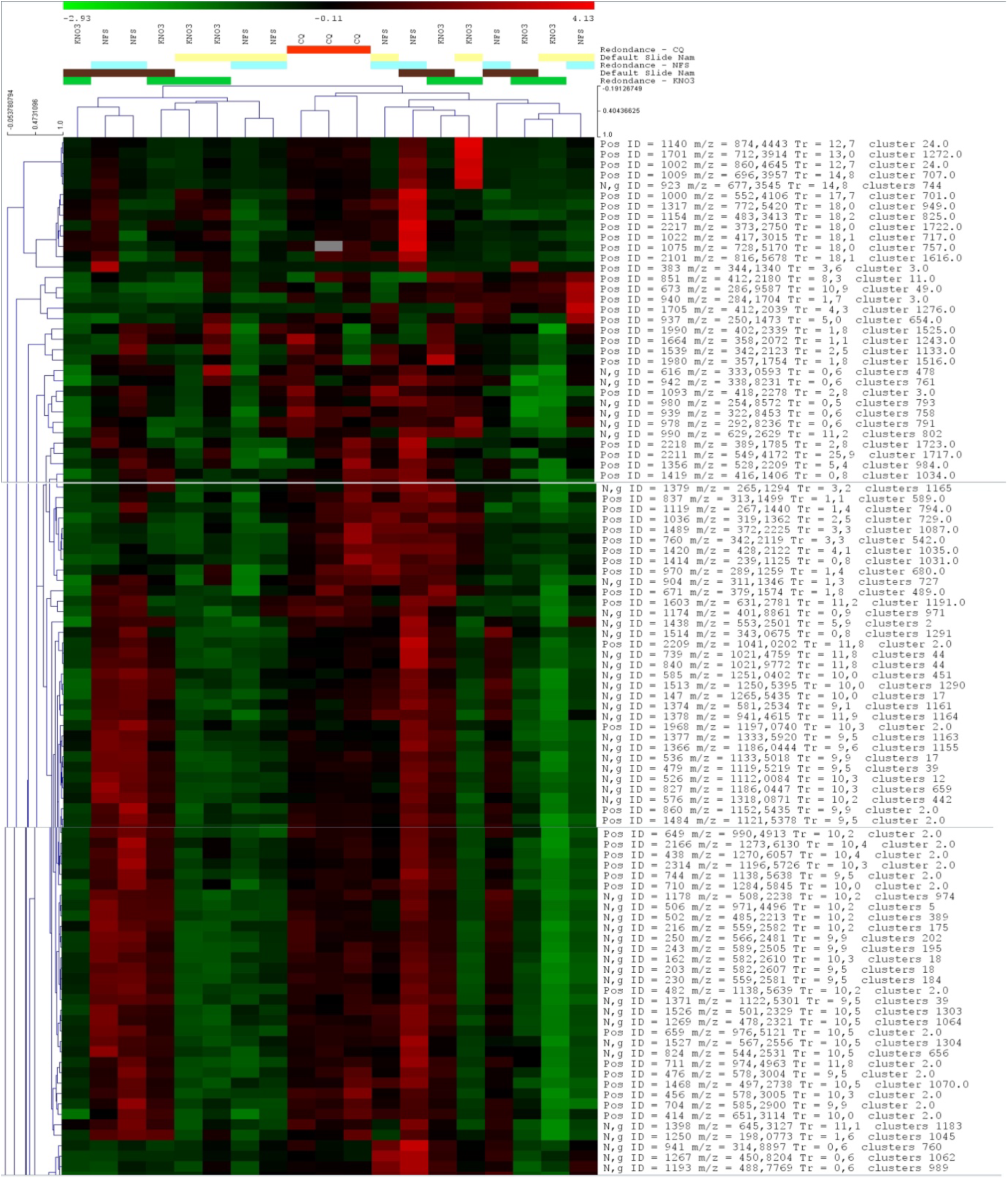
Unsupervised clustering of LC-MS compounds. Heatmap showing the top 100 features from LC-MS analysis.

**Fig. S7:**
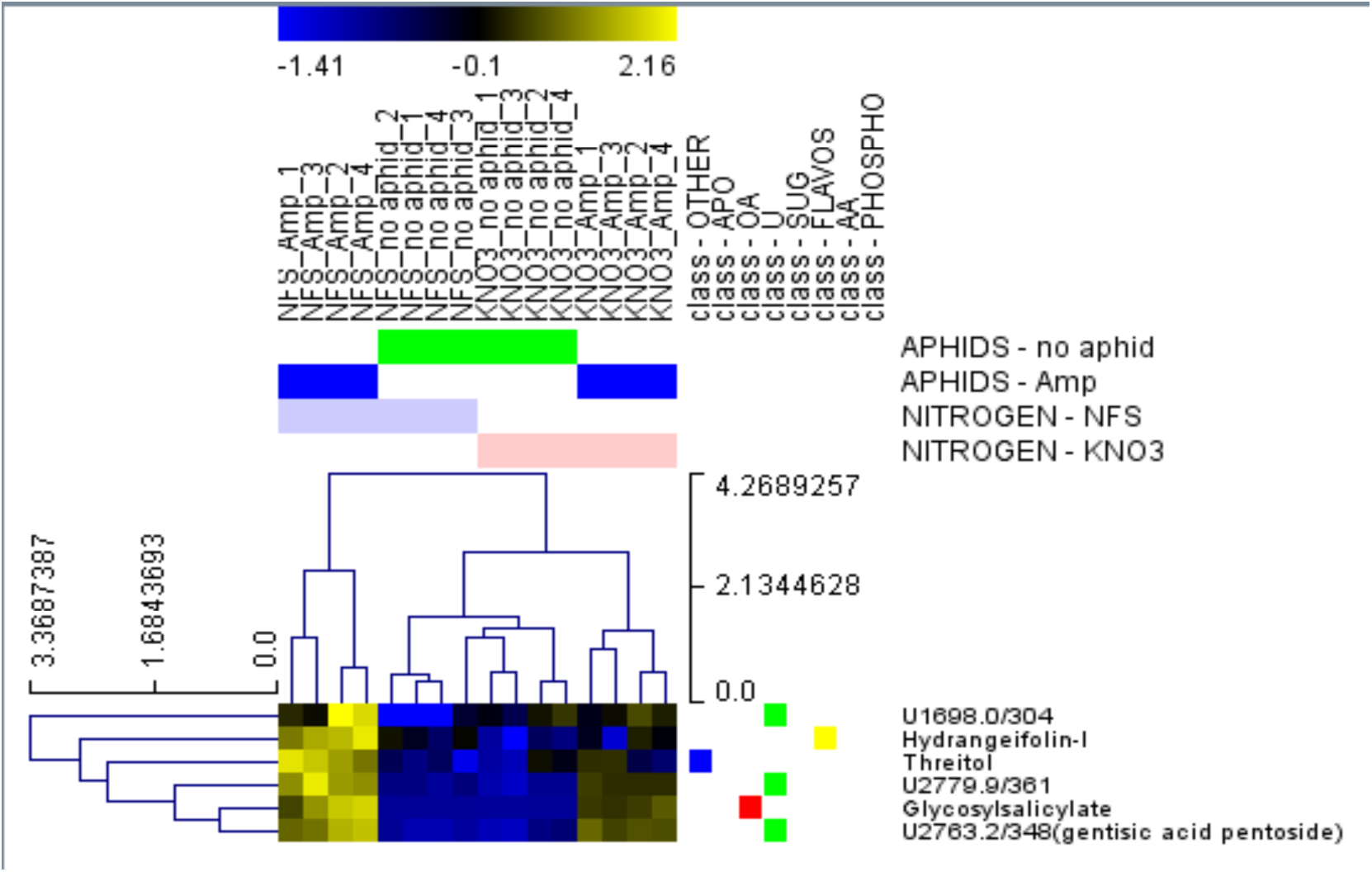
NFS and Aphid infestation significantly modifies the leaf metabolite profile. Heatmap of 2-way Anova analysis from GC-MS showing unsupervised hierarchical clustering of compounds separated by the interaction of NFS and aphid infestation. NFS = Nitrogen fixing symbiosis, KNO3 = Non inoculated, Amp = aphid infested, no aphids = Control.

**Table S1:**
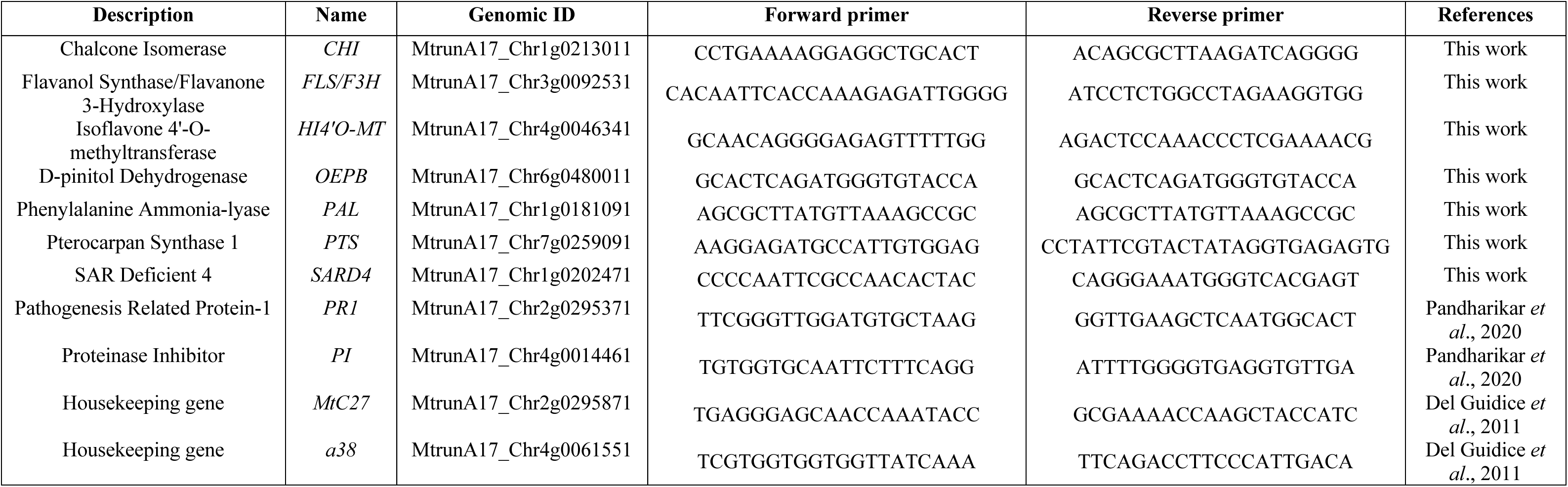
List of genes and oligos sequences used for RT-qPCR analysis.

