## Supplementary material for "Differential induction of *Medicago truncatula* defence metabolites in response to rhizobial symbiosis and pea aphid infestation": Benjamin et al._Supplementary data

*Both authors supervised this work.

**Supplementary Data**

Datasets available as: Benjamin, Goodluck, 2024, "Supplementary Data for Differential induction of Medicago truncatula defence metabolites in response to rhizobial symbiosis and pea aphid infestation", <https://doi.org/10.57745/PCDG8H>, Recherche Data Gouv, V1, UNF:6:A0SbGGbxQocqa6JsmMuoMA== [fileUNF]

Dataset S1: File containing qPCR Data (qPCR CT Data.xlsx)

Dataset S2: File containing GC-MS and LC-MS metabolomics data (Metabolomics_data.xlsx)

Dataset S3: File containing Significant accumulated metabolites analysed by Anova and Venn Analysis (Significant metabolites_Venn.xlsx)

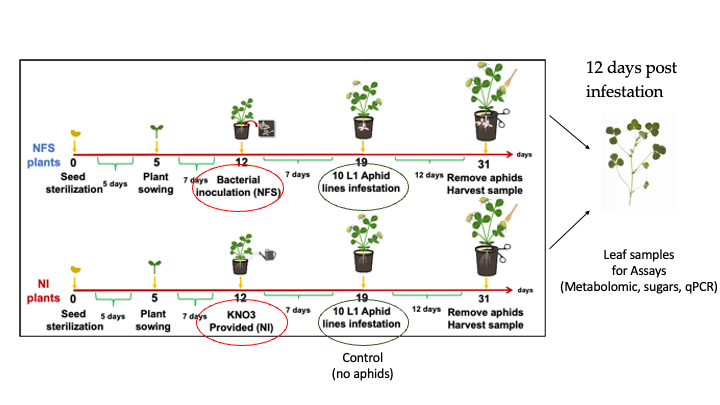

**Fig. S1: Experimental setup.** Diagram showing timeline leading up to assay with key date points for bacteria inoculation and aphid infestation marked up. Each pot contained six plants and 4 pots were made for each condition per biological replicate. Assay was performed using 4 biological replicates produced using the same timeline.

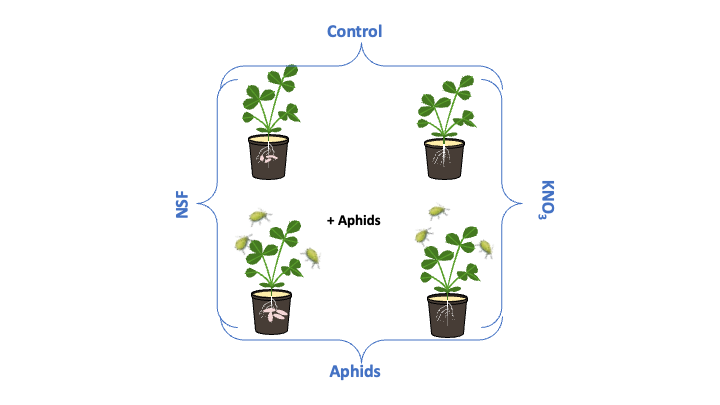

**Fig. S2: Experiment treatment classification.** Diagram showing the classification of studied treatments into 4 groups, NFS_Control, NFS_Amp, NI_Control (KNO_3_) and NI_Amp (KNO_3_).

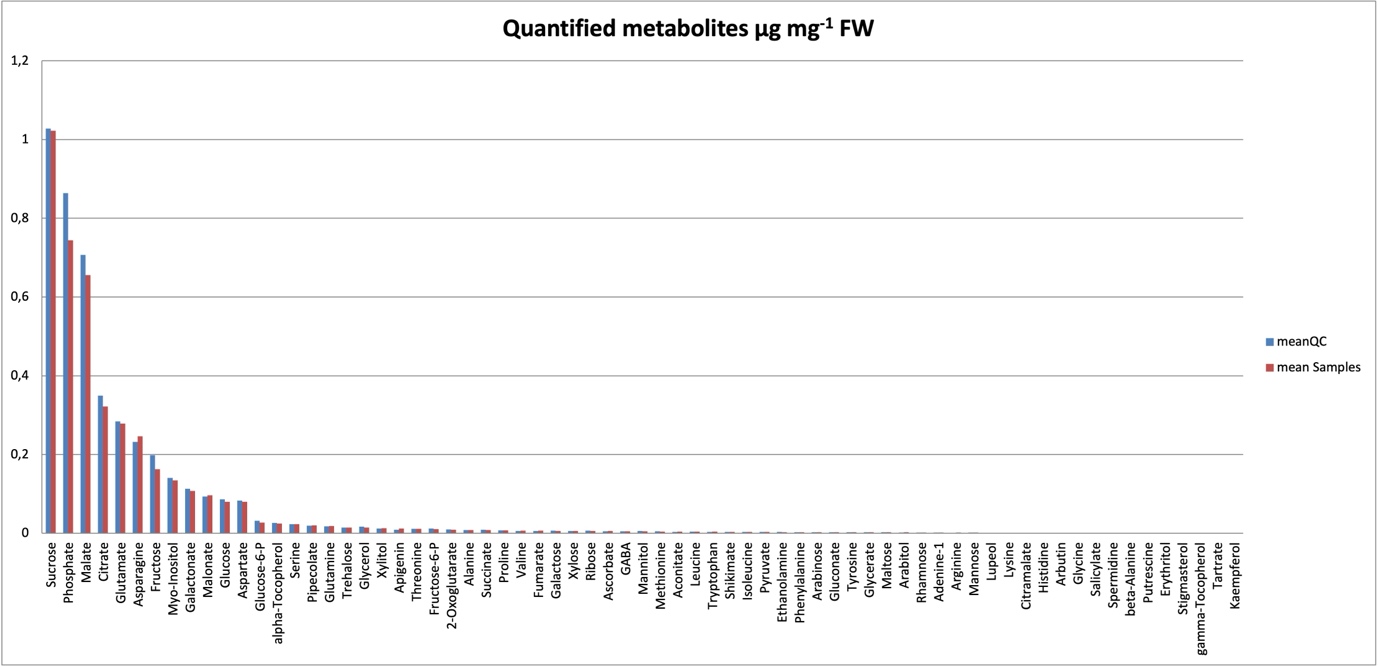

**Fig. S3: Abundance of quantified metabolites.** Graph of quantified metabolites from GC-MS analysis showing accuracy of abundance by comparing the mean obtained from the quality controls (QC, blue bars) and mean obtained from the samples (Samples, red bars).

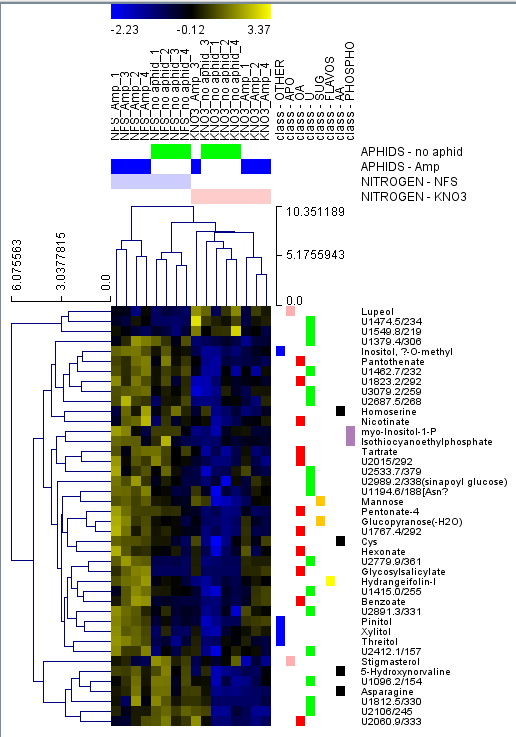

**Fig. S4: Nitrogen source significantly modifies the leaf metabolite profile.** Heatmap of 2-way Anova analysis from GC-MS showing unsupervised hierarchical clustering of compounds separated by nitrogen source. NFS = Nitrogen fixing symbiosis, KNO3 = Non inoculated, Amp = aphid infested, no aphids = Control.

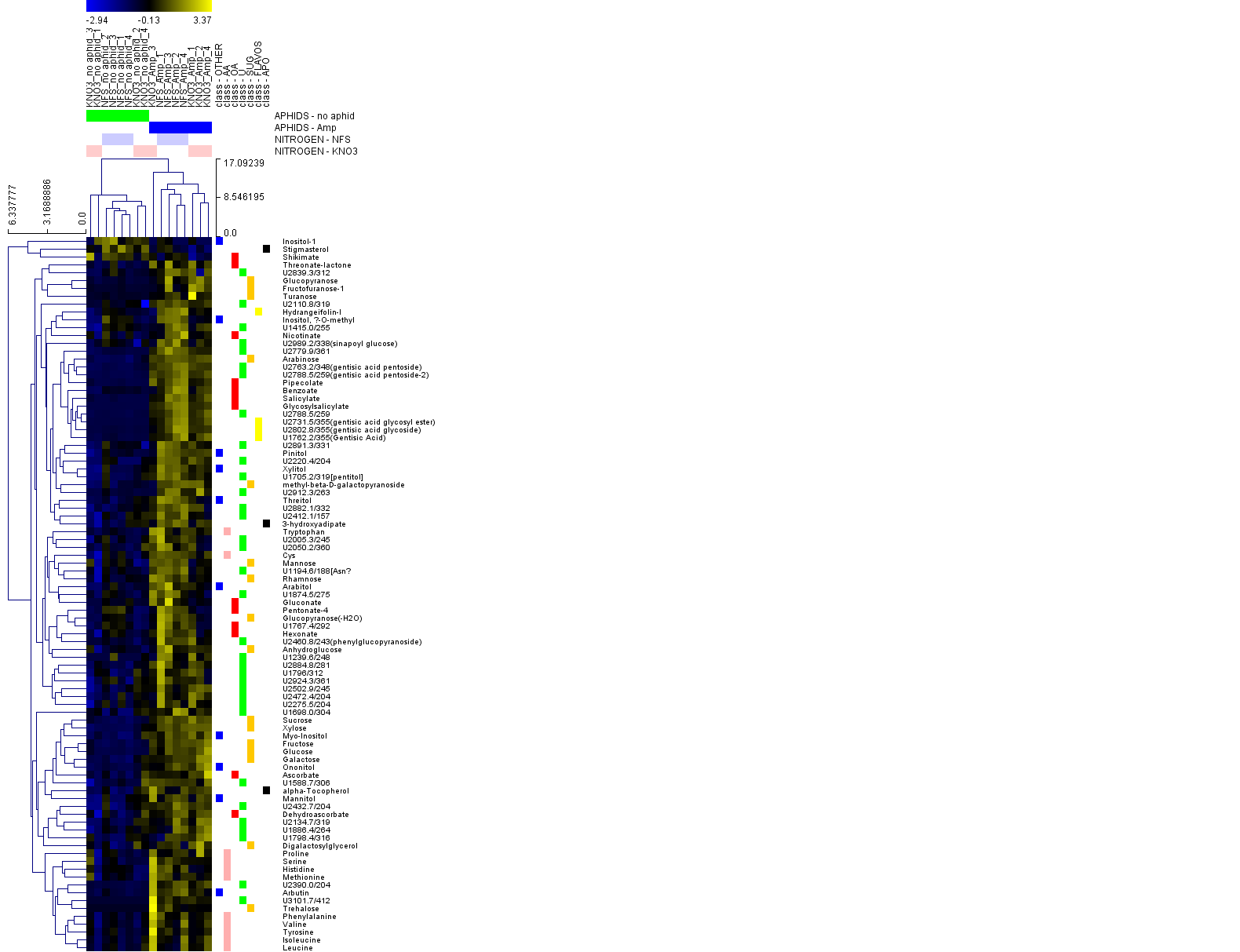

**Fig. S5: Aphid infestation significantly modifies the leaf metabolite profile.** Heatmap of 2-way Anova analysis from GC-MS showing unsupervised hierarchical clustering of compounds separated by aphid infestation. NFS = Nitrogen fixing symbiosis, KNO3 = Non inoculated, Amp = aphid infested, no aphids = Control.

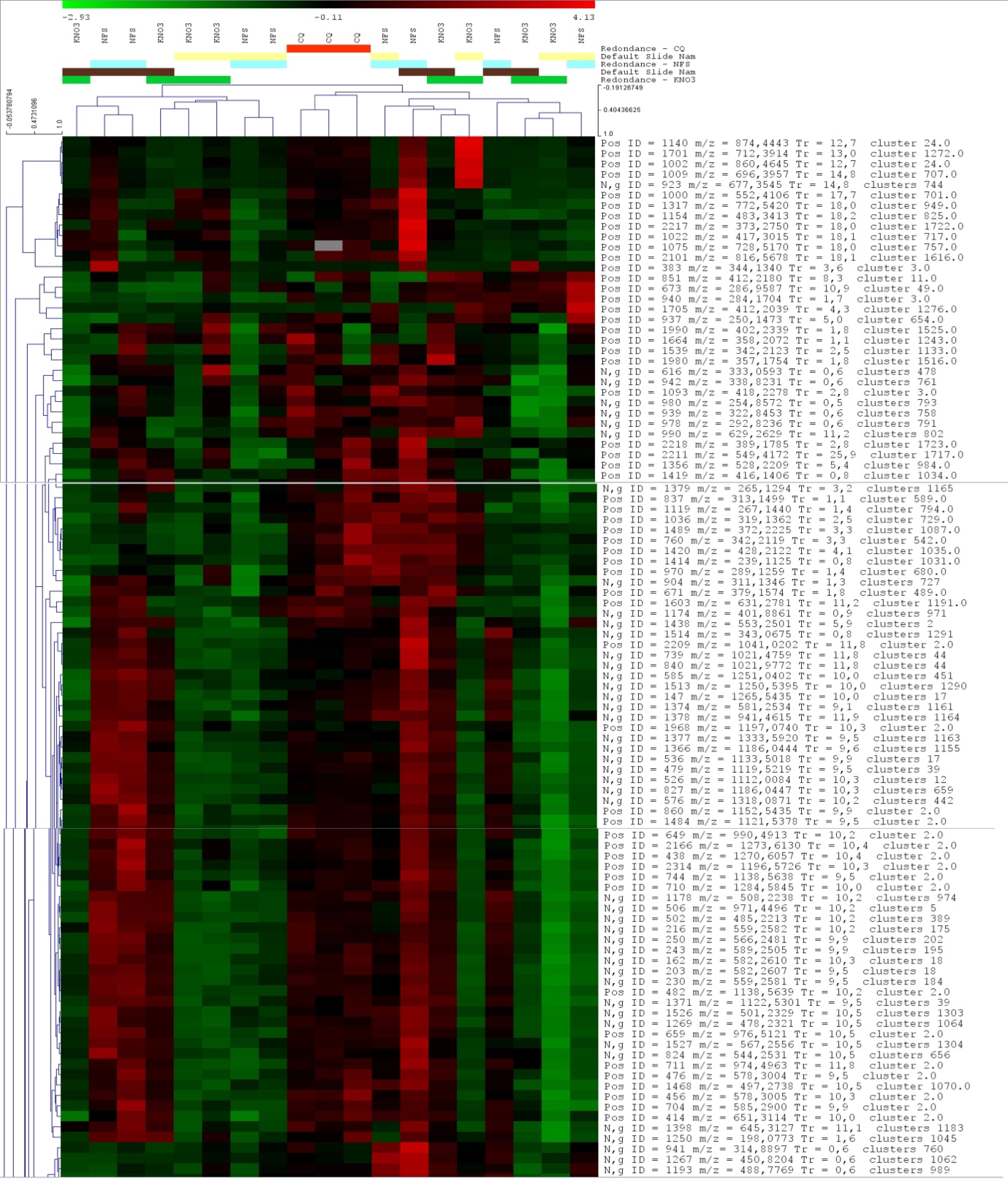

**Fig. S6: Unsupervised clustering of LC-MS compounds.** Heatmap showing the top 100 features from LC-MS analysis.

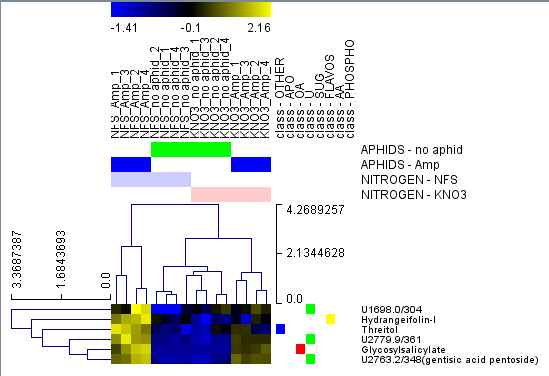

**Fig. S7: NFS and Aphid infestation significantly modifies the leaf metabolite profile.** Heatmap of 2-way Anova analysis from GC-MS showing unsupervised hierarchical clustering of compounds separated by the interaction of NFS and aphid infestation. NFS = Nitrogen fixing symbiosis, KNO3 = Non inoculated, Amp = aphid infested, no aphids = Control.

**Table S1: List of genes and oligos sequences used for RT-qPCR analysis.**

| **Description** | **Name** | **Genomic ID** | **Forward primer** | **Reverse primer** | **References** |
| --- | --- | --- | --- | --- | --- |
| Chalcone Isomerase | *CHI* | MtrunA17_Chr1g0213011 | CCTGAAAAGGAGGCTGCACT | ACAGCGCTTAAGATCAGGGG | This work |
| Flavanol Synthase/Flavanone 3-Hydroxylase | *FLS/F3H* | MtrunA17_Chr3g0092531 | CACAATTCACCAAAGAGATTGGGG | ATCCTCTGGCCTAGAAGGTGG | This work |
| Isoflavone 4'-O-methyltransferase | *HI4'O-MT* | MtrunA17_Chr4g0046341 | GCAACAGGGGAGAGTTTTTGG | AGACTCCAAACCCTCGAAAACG | This work |
| D-pinitol Dehydrogenase | *OEPB* | MtrunA17_Chr6g0480011 | GCACTCAGATGGGTGTACCA | GCACTCAGATGGGTGTACCA | This work |
| Phenylalanine Ammonia-lyase | *PAL* | MtrunA17_Chr1g0181091 | AGCGCTTATGTTAAAGCCGC | AGCGCTTATGTTAAAGCCGC | This work |
| Pterocarpan Synthase 1 | *PTS* | MtrunA17_Chr7g0259091 | AAGGAGATGCCATTGTGGAG | CCTATTCGTACTATAGGTGAGAGTG | This work |
| SAR Deficient 4 | *SARD4* | MtrunA17_Chr1g0202471 | CCCCAATTCGCCAACACTAC | CAGGGAAATGGGTCACGAGT | This work |
| Pathogenesis Related Protein-1 | *PR1* | MtrunA17_Chr2g0295371 | TTCGGGTTGGATGTGCTAAG | GGTTGAAGCTCAATGGCACT | Pandharikar *et al*., 2020 |
| Proteinase Inhibitor | *PI* | MtrunA17_Chr4g0014461 | TGTGGTGCAATTCTTTCAGG | ATTTTGGGGTGAGGTGTTGA | Pandharikar *et al*., 2020 |
| Housekeeping gene | *MtC27* | MtrunA17_Chr2g0295871 | TGAGGGAGCAACCAAATACC | GCGAAAACCAAGCTACCATC | Del Guidice *et al*., 2011 |
| Housekeeping gene | *a38* | MtrunA17_Chr4g0061551 | TCGTGGTGGTGGTTATCAAA | TTCAGACCTTCCCATTGACA | Del Guidice *et al*., 2011 |
